# Detecting Boolean Asymmetric Relationships with a Loop Counting Technique and its Implications for Analyzing Heterogeneity within Gene Expression Datasets

**DOI:** 10.1101/2022.08.04.502792

**Authors:** Haosheng Zhou, Wei Lin, Sergio R. Labra, Stuart A. Lipton, Jeremy A. Elman, Nicholas J. Schork, Aaditya V. Rangan

## Abstract

Many traditional methods for analyzing gene-gene relationships focus on positive and negative correlations, both of which are a kind of ‘symmetric’ relationship. Biclustering is one such technique that typically searches for subsets of genes exhibiting correlated expression among a subset of samples. However, genes can also exhibit ‘asymmetric’ relationships, such as ‘if-then’ relationships used in boolean circuits. In this paper we develop a very general method that can be used to detect biclusters within gene-expression data that involve subsets of genes which are enriched for these ‘boolean-asymmetric’ relationships (BARs). These BAR-biclusters can correspond to heterogeneity that is driven by asymmetric gene-gene interactions, e.g., reflecting regulatory effects of one gene on another, rather than more standard symmetric interactions. Unlike typical approaches that search for BARs across the entire population, BAR-biclusters can detect asymmetric interactions that only occur among a subset of samples. We apply our method to a single-cell RNA-sequencing data-set, demonstrating that the statistically-significant BAR-biclusters indeed contain additional information not present within the more traditional ‘boolean-symmetric’-biclusters. For example, the BAR-biclusters involve different subsets of cells, and highlight different gene-pathways within the data-set. Moreover, by combining the boolean-asymmetric- and boolean-symmetric-signals, one can build linear classifiers which outperform those built using only traditional boolean-symmetric signals.

## 1. Introduction

IT is widely recognized that relationships between genes play an important role in biological research [1]. Generally speaking, most common methods for finding/identifying genetic relationships focus on positive and negative correlations, which we’ll refer to below as ***symmetric relationships*** [2]–[4]. In this paper, however, we’ll focus on ***asymmetric relationships***, which are a different kind of genetic relationship introduced and studied in [5]–[9]. Our main goal is to introduce a version of ‘biclustering’ – a form of heterogeneity analysis typically defined in terms of symmetric relationships – to also include these asymmetric relationships. Since there is growing evidence that a significant fraction of biologically-relevant gene-gene interactions are indeed asymmetric [6], our ultimate goal is to provide a simple preprocessing strategy that can be used to extend current biclustering tools to capture these asymmetric interactions.

We begin by reviewing some of the properties of asymmetric relationships, focusing specifically on ‘boolean-asymmetric-relationships’ (BARs) and briefly summarizing some recent research highlighting the importance of these asymmetric-relationships in genetic data. We then describe the notion of a ‘bicluster’ as it applies to BARs: in this context a bicluster might comprise a subsets of samples which exhibit certain asymmetric-relationships across a subset of genes; relationships that are not shared by the rest of the samples or genes within the data. As a first step towards finding these ‘BAR-biclusters’, a very simple strategy for reorganizing data is introduced so that BARs within the original data-set are represented as boolean-symmetric-relationships (i.e., BSRs) within the transformed data-set. This data-reorganization then allows standard strategies for analyzing symmetric-relationships to be used as tools for analyzing asymmetric relationships. In particular, standard strategies for finding traditional BSR-biclusters can be used on a transformed data-set to find BAR-biclusters. We illustrate this technique using a single-cell RNA-sequencing data-set, demonstrating that the BAR-biclusters are not only strongly statistically-significant in their own right, but contain information that is distinct from the more traditional BSR-biclusters. Given these results, we expect that BAR-biclustering can be a useful complement to BSR-biclustering, allowing for a new kind of heterogeneity analysis.

### A. Reviewing Boolean Asymmetric Relationships (BARs)

Traditional analytical tools typically look for relationships between genes and/or samples which involve positive- or negative-correlations. These relationships have the property that if gene *X* is positively correlated with gene *Y*, then the reverse is also true (i.e., *Y* is correlated with *X*). For this reason these correlations are sometimes referred to as ‘symmetric’ relationships. A more general kind of interaction can involve ‘asymmetric’ relationships; knowledge of gene *X* may strongly inform the value of *Y*, without the reverse holding true. Interest in asymmetric-relationships has grown in recent years, and several informative asymmetric-relationships have been discovered in a variety of cellular- and genetic-data-sets, e.g., in the context of cancer data-sets [7], [8], [12], [13], [23], [38]–[40], hypoxia-related genes [34], and cell proliferation [9]. For instance, a boolean analysis in [9] identified more than two dozen new genes involved in cell cycle processes, with their effects subsequently validated using a recent RNA-sequencing data-set. As one example, the gene CCNB1, which encodes cyclin B1, is ‘gated’ by the gene CASC5, a cell-cycle gene, meaning that the expression level of CCNB1 can only be high if CASC5 is expressed at a high level.

For the purpose of illustration, we’ll use the terminology from [5]. Consider two random variables *X* and *Y*, each with *M* observations *x*_*m*_ ∈ ℝ and *y*_*m*_ ∈ ℝ, respectively. Each one of the observed points (*x*_*m*_, *y*_*m*_) ∈ ℝ^2^ lies in one of the four quadrants of the plane. A typical ‘symmetric’ relationship between *X* and *Y* is defined as either an abundance of observed points in the first and third quadrants (i.e., positive correlation) or as an abundance of points in the second and fourth quadrants (i.e., negative correlation). Thus, if we denote the joint probability density of *X* and *Y* as *ρ*(*x, y*), a symmetric relationship involves an elevation of *ρ*(*x, y*) in opposite quadrants. By contrast, an ‘asymmetric’ relationship refers to any nonuniformity in *ρ*(*x, y*) which is not inversion-symmetric, e.g., if *ρ*(*x, y*) is elevated in only one of the four quadrants.

While there are many different kinds of asymmetric relationships, only certain asymmetric relationships can persist after centering the data (i.e., centralizing the marginal distributions for both *X* and *Y* with respect to the means). Assuming the data is centered, an asymmetric relationship must involve an elevation of *ρ*(*x, y*) in three quadrants (i.e., *ρ*(*x, y*) can have only exactly one relatively vacant quadrant). An illustration of this kind of asymmetric relationship is shown in figure 1.

**Fig. 1.**
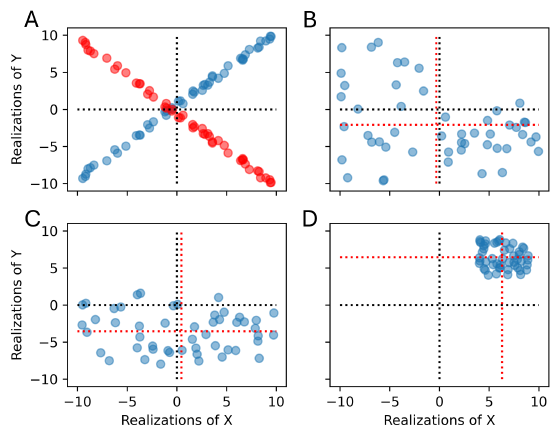
Symmetric and asymmetric relationships. (A) a positive correlation (blue) and a negative correlation (red) as symmetric relationships. (B) the kind of asymmetric relationships we are focusing on, with few points in the first quadrant. (C) and (D) asymmetric relationships under the standard coordinate axes (black dotted lines). However, they are no longer asymmetric if the points are centered with respect to the marginal means (red dotted lines).

In this paper we focus on a particular kind of asymmetric relationship: a ‘boolean-asymmetric-relationship’ (BAR), which has proven to be of interest for genetic analysis [5], [6]. In this kind of relationship we assume that the random variables *X* and *Y* are categorical variables that can each take on up to two values. This can happen when *X* and *Y* are naturally binary, or with continuous variables that have been binarized based on a given threshold [5]. As an example, consider *X* to be an indicator of a person’s sex and *Y* to be an indicator of balding. The joint distribution *ρ*(*x, y*) is almost limited to three quadrants, as *y*_*m*_ is rarely positive if *x*_*m*_ is female.

Since each BAR involves underexpression in a single quadrant, there are four different kinds of BARs:

- *Y* ⇒ *X*: This direction-specific interaction implies that *X* is necessary for *Y* . Put another way, *Y* cannot be high unless *X* is high (i.e., the 2^*nd*^ quadrant is relatively empty).
- *X* ⇒ *Y* : This is the converse of the previous interaction (i.e., the 4^*th*^ quadrant is relatively empty).
- *X* ‘or’ *Y* : This two-way interaction implies that *either X or Y* is on (i.e, the 3^*rd*^ quadrant is relatively empty).
- *X* ‘nand’ *Y* : This two-way interaction implies that *either X or Y* is off (i.e, the 1^*st*^ quadrant is relatively empty).

Note that the ‘*Y* ⇒ *X*’ and ‘*X* ⇒ *Y* ‘ relationships will reverse their directions if the *x* and *y* labels are interchanged; these two relationships are ‘directed’ in the sense that the ordering of *x* and *y* matters. Meanwhile, the ‘or’ and ‘nand’ relationships do not change if the *x* and *y* labels are interchanged; these two relationships are ‘undirected’.

### B. Prior work demonstrating the importance of BARs

Boolean-asymmetric-relationships (BARs, described above) are relevant in many biological contexts, and have been found present in many different gene-expression data-sets [5]–[9], [12], [13], [23], [34]–[40]. As alluded to above, the genegene BARs each suggest an associated genetic interaction; one gene can be necessary for the other, or the two genes can be linked in an ‘or’ or ‘nand’ pair. In this fashion, BARs that are associated with, e.g. a disease-condition, can be interpreted as playing a role in a larger gene-interaction-network, in which the individual BARs suggest potential causal interactions between genes that are responsible for the disease.

These benefits suggest that BARs might serve as an important supplement to more traditional correlation-based analyses (i.e., involving BSRs) when analyzing genetic data. Indeed, recent research corroborates this perspective. Several statistically-significant BARs have been observed relating certain genes to the developmental stages of tumors sampled in colon, bladder and prostate cancers [38]–[40]. Along similar lines, significant BARs have been found associated with cell-differentiation, cellular-development and tissue-function [9], [34]–[37]. More recent work has shown that, while many BARs differentiate between different cell-types in the microbiome, there are also many BARs that are strongly conserved across microorganisms, potentially highlighting ‘universal’ gene-functions important for survival [6].

The results above motivate our interest in finding BARs and our study of the Alzheimer’s hiPSC-derived organoids from [25].

### C. Prior computational strategies for identifying BARs

Due to the importance and ubiquity of BARs, many different methods have been developed to identify and characterize BARs within genetic data. To list a few of them, the StepMiner algorithm [10] is combined with a sparsity test to efficiently find statistically-significant BARs, while its generalized version [11] finds generalized BARs corresponding to three-way-interactions between genes. The StepMiner algorithm is also used as a component in the BECC algorithm [36] which forms scores indicative of boolean-equivalence between gene probesets. Other work includes the MiDReG method [35] which uses significant BARs to identify developmentally regulated genes. Related work includes [16] which investigates the choice of binarization threshold for defining BARs, and [13] which investigates the conditional probability of overexpression (i.e., asymmetric-relationships) between pairs of genes.

The abundance of asymmetric-relationships found in genetic data-sets has motivated a variety of strategies for constructing ‘Boolean-networks’ – connected graphs with nodes corresponding to genes and edges corresponding to interactions between genes (e.g., BARs). Earlier work by [32] provides a flexible method for constructing ‘probabilistic’ Boolean-networks which can (in principle) involve multiple boolean functions when predicting any particular gene’s expression. Almost concurrently, work by [33] focuses on finding families of Boolean-networks consistent with a given data-set. More recent work by [31] draws parallels between Boolean-networks and Markov chains, while [15] investigates the dynamic aspects of Boolean-networks which incorporate time-delays. Boolean-networks have been analyzed in the context of lungcancer risk [12] and yeast cell-cycle data [14].

### D. Our Motivation and Contribution

As far as we are aware, most methods for identifying BARs focus on detecting statistically-significant relationships between small groups of genes, with the majority focusing on pairs of genes at a time. These methods typically estimate the strength of any particular gene-gene BAR by considering all the samples available, highlighting the pairwise gene-gene-BARs with the strongest evidence (e.g., the highest likelihood calculated across the samples). Our main goal in this paper is to introduce a different paradigm for delineating BARs – that of a ‘BAR-bicluster’ – which allows for BARs that extend across larger subsets of genes, but which can be limited in scope to only a subset of the samples.

To describe this notion in more detail, we first introduce some notation, with terminology taken from single-cell RNA-sequencing (scRNA-seq). Consider a data-set *D* including *M* samples (e.g., cells isolated in a scRNA-seq experiment), each observed across *N* variables (e.g., unique-molecular-identifiers, which we’ll refer to as ‘genes’). The value of the *n*^th^-gene observed within the *m*^th^-cell is stored at the array location *D*_*mn*_.

As mentioned above, many of the existing analytical methods for detecting BSRs and/or BARs in genetic data emphasize relationships between gene-pairs that are statistically significant when considered across the entire sample-population. That is to say, they focus on identifying which of the 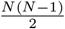 individual gene-pairs are significant when considered across all *M* of the cells. These all-cell gene-gene relationships are certainly important, but are not the only important relationships that exist. Other important relationships include ‘clusters’ and ‘biclusters’.

A gene-cluster is a subset of genes among which each pair of genes may only be weakly linked but jointly exhibit a collective signal (across all the cell-samples) that is strongly significant. There are many different kinds of BSR-clustering algorithms [41], [42], and some of the methods referenced above can be used to identify clusters of genes that share the same kinds of BARs [36].

On the other hand, a ***bicluster*** is defined as a subset of genes which only exhibits a collective signal across a subset of cells. Importantly, the relationships that exist within a bicluster (i.e., between the genes and/or between cells) typically do *not* exist within the rest of the data-array. Thus, a bicluster is a submatrix (defined in terms of a row-and column-subset) of the data-array *D* which exhibits a strongly statistically-significant structure which is typically only weakly significant (or even insignificant) when measured across the entire dataarray.

Biclusters can highlight the drivers of heterogeneity within a data-set, which can be useful for developing new kinds of classifications, such as defining new cell-subtypes [21], [22], [24]. There is an abundance of literature discussing the kinds of biclusters which manifest in gene-expression data-sets, as well as various strategies for finding them [17], [43], [44]. In the discussion below we’ll use both terms ‘cluster’ and ‘bicluster’ to refer to biclusters.

Generally speaking, detecting all the statistically-significant biclusters is very difficult, mainly because the number of possible biclusters is exponential in the number of cells and genes (i.e., there are 2^*M*+*N*^ different biclusters one could consider). Nevertheless, there are several biclustering methods which perform well in certain regimes [18]–[24]. Typically, these methods are built using strategies which either implicitly or explicitly search for BSRs between genes. Thus, the commonly available packages for biclustering gene expression data are designed to identify biclusters which exhibit an abundance of BSRs (i.e., ‘BSR-biclusters’). See [19] for the Louvain Clustering algorithm, [20] for the UMAP method and see [21]–[24] for other methods which perform BSR-biclustering.

In the remainder of this paper we introduce a simple strategy for finding biclusters which exhibit an abundance of BARs (i.e., ‘BAR-biclusters’). These BAR-biclusters can reveal heterogeneity which is distinct from that associated with BSR-biclusters. It is important to note that BAR-biclusters include both BARs and standard BSRs, which can be differentiated with a post-processing step presented in III-G. However, it is worth clarifying that in most practical scenarios, BARs and BSRs cannot be completely disentangled. A collection of genes that are enriched for BARs will often contain a subset of genes within it that are boolean-equivalent, thus expressing BSRs. Our strategy for finding BAR-biclusters boils down to a simple preprocessing step, where the original data is split to form a slightly larger dataset, which then can be passed to any state-of-the-art biclustering algorithm that looks for BSR-biclusters. In other words, **our contribution in this paper is not an entirely distinct methodology but rather a general strategy that turns any BSR-bicluster finder into a BAR-bicluster finder**.

As a proof of principle, we illustrate our strategy using the BSR-biclustering algorithm published in [18]. We combine this BSR-biclustering algorithm – called ‘loop-counting’ in [18] – with our preprocessing step, referring to this composition as the ‘loop-counting-asymmetric’ (LCA) algorithm. We then use this LCA algorithm to find BAR-biclusters within an organoid dataset [25]. Using our new method, we demonstrate that BAR-biclusters contain information which can complement traditional BSR-biclustering.

### E. The Organoid Dataset

As mentioned above, we will apply our LCA algorithm to an scRNA-seq data-set studied in [25]. This data-set involves multiple cells sampled from organoids grown from patientderived human induced pluripotent stem cells (hiPSCs). These organoids were designed to study Alzheimer’s disease, and were prepared with three different isogenic types (for details see [25], [53], [54]). The first is the baseline wild-type (WT), the second involves a mutation in the amyloid-precursor-protein (APP), and the third a mutation in presenilin-1 (PS1); both mutations are strongly associated with inherited alzheimer’s disease. The organoids were allowed to develop (under conditions described in [25]) and were sampled at three different stages in development: 6-weeks, 3-months and 6-months. The cells sampled from the organoids were analyzed using scRNA-seq, and classified into cell-types by a supervised classifier trained on previous annotations (see [26]–[28]). Our analysis below will search for BAR-biclusters within the cell-types, identifying subsets of cells which exhibit a statistically-significant abundance of BARs as well as BSRs across subsets of genes. As we’ll show below, these BAR-biclusters contain information which is distinct from the typical BSR-biclusters found within the same data-set. As we’ll discuss later on, we believe this extra information can be useful for basic tasks such as classification and identification, as well as for understanding the underlying genetic drivers for the heterogeneity within the samples.

## II. Materials and methods

### A. Column Splitting Technique

The main computational technique we will use in this paper is ‘column-splitting’ (described below), which can transform any BAR-biclustering problem into a BSR-biclustering problem.

Consider the data-array *D* ∈ ℝ^*M×N*^ mentioned above. Let’s assume, in keeping with [5], that some thresholds have already been chosen to convert each entry of *D* from gene-expression values into categorical labels. Thus, each entry *D*_*mn*_ is either “low” (under-expressing), “high” (over-expressing) or “intermediate” (not significantly expressing, typically neglected). To describe our column-splitting technique, we’ll use ‘matlab’ notation throughout the paper: referring to the *n*^th^-column of *D* as *D*_:,*n*_, and the *m*^th^-row of *D* as *D*_*m*,:_.

Our column splitting technique proceeds as follows: Split each column of *D* into two columns to get a new binarized data-array *B* ∈ ℝ^*M×*2*N*^ . Each column *D*_:,*n*_ from *D* will correspond to the two columns *B*_:,2*n*−1_ and *B*_:,2*n*_ in *B*. If *D*_*mn*_ is low, then *B*_*m*,2*n*−1_ = 1 and *B*_*m*,2*n*_ = 0. If *D*_*mn*_ is high, then *B*_*m*,2*n*−1_ = 0 and *B*_*m*,2*n*_ = 1. If *D*_*mn*_ is intermediate, then *B*_*m*,2*n*−1_ = 0 and *B*_*m*,2*n*_ = 0. Thus, each column of *D* is split into one column indicating its ‘low’ state and one column indicating its ‘high’ state.

The technique is simple but effective. Similar data-splitting techniques have been used in other applications [55], but here we focus on how this column-splitting facilitates the detection of BARs. To illustrate how column-splitting works in this context, assume that *D* has only two genes, indexed by columns *n* = 1 and *n* = 2, respectively. Furthermore, let’s assume that genes 1 and 2 exactly satisfy an ‘nand’-type BAR, implying that *D*_*m*1_ and *D*_*m*2_ each takes on a range of labels, but cannot both be high simultaneously. Thus, the columns *B*_:,2_ and *B*_:,4_ (corresponding to the high-states of *D*_:,1_ and *D*_:,2_, respectively) must have a zero dot-product, while the other column-pairs (*B*_:,1_, *B*_:,3_), (*B*_:,1_, *B*_:,4_) and (*B*_:,2_, *B*_:,3_) can all have nonzero dot-products. Thus, when contrasted against a scenario where genes 1 and 2 are independent random variables, the column-pair (*B*_:,2_, *B*_:,4_) exhibits a lower-than-expected correlation. Similarly, the column-pairs (*B*_:,1_, *B*_:,3_), (*B*_:,1_, *B*_:,4_) and (*B*_:,2_, *B*_:,3_) all exhibit a higher-than-expected correlation. A standard measurement of BSR can be used to detect these correlations amongst the columns of *B*, thus highlighting the BAR between the columns of *D*.

A similar statement holds more generally. Given any two distinct columns of *D*, say *D*_:,*n*_ and *D*_:,*n′*_, the corresponding columns of *B* form 4 nontrivial column-pairs: (*B*_:,2*n*−1_, *B*_:,2*n*′−1_), (*B*_:,2*n*−1_, *B*_:,2*n*′_), (*B*_:,2*n*_, *B*_:,2*n*′−1_) and (*B*_:,2*n*_, *B*_:,2*n′*_) (Note that (*B*_:,2*n*−1_, *B*_:,2*n*_) and (*B*_:,2*n′*−1;_, *B*_:,2*n′*_) are orthogonal by construction). Each of the nontrivial column-pairs in *B* corresponds to a quadrant regarding the original columns of *D*. Each BAR corresponds to underexpression within a single quadrant. Thus, if the column-pair (*D*_:,*n*_, *D*_:,*n′*_) within *D* exhibits a BAR, then three of the the corresponding column-pairs within *B* will exhibit an elevated dot-product (i.e., a positively-correlated BSR) while the remaining quadrant will be underexpressed.

To sum up, the separation of ”low” and ”high” states turns the problem of finding BAR-biclusters into a problem of finding BSR-biclusters in a slightly larger matrix *B*. In principle, the matrix *B* can be processed by any algorithm which can detect BSR-biclusters; below we will use the specific BSR-biclustering algorithm in [18]. We also remark that, by splitting columns, the BSR-biclusters of *B* can indicate both BARs and BSRs within *D*. For this reason we will perform BSR-biclustering and BAR-biclustering simultaneously in our numerical experiments below; by comparing the results we will be able to isolate BAR-specific signals.

### B. Loop Counting Low Rank (LCLR)

As stated above, the column splitting technique is a very simple strategy for finding BAR-biclusters. This strategy can make use of any method which finds BSR-biclusters, but for demonstration we make use of the LCLR method for finding BSR-biclusters. This method is described in [18], and we review the important aspects here.

The LCLR method provides an approximate solution to the problem of finding submatrices with low numerical rank within a large matrix *D* ∈ ℝ^*M×N*^ .

Given any binary matrix *B* ∈ ℝ^*M×N*^ as a binarized version of *D*, with its entries taking value ±1, the LCLR method accumulates information about ‘loops’, referring to 2 × 2 submatrices within the binarized-array *B*. Because each entry *B*_*mn*_ is binary, there are only 2^4^ = 16 possibilities for each loop, and the ranks (i.e., dimensions) of each loop can be efficiently calculated and accumulated to form row- and column-scores.

The row-score for any row *m* is the number of rank-1 loops containing that row minus the number of rank-2 loops containing that row. This particular definition of the row-score is why LCLR is called a ‘loop-counting’ algorithm. The rank of any loop is determined by the product of its entries.

For example, given a single loop of a binary matrix:

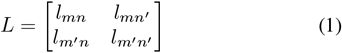

The product *l*_*mn*_*l*_*mn′*_ *l*_*m′n*_*l*_*m′n′*_ is +1 if and only if the loop is of rank-1, and the product is −1 otherwise. Each loop of rank-1 contributes +1 to the score of the *i*-th row, while each loop of rank-2 contributes −1, consistent with the definition above for the score of a row.

By simple linear algebra, the score of the *m*^*th*^ row can be written as:

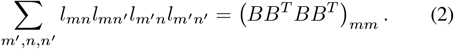

Similarly, the score of the *n*^*th*^ column can be written as:

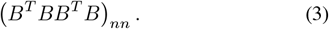

The LCLR algorithm proceeds iteratively: calculating the row- and column-scores, eliminating rows and columns with low scores, and then recalculating the scores and eliminating more rows and columns until all rows and columns are eliminated. Those rows and columns which are retained the longest are highly likely to be part of a low-rank bicluster within the original data-set, provided that a bicluster of sufficient signal-strength exists (see Appendix A for more details, and [18] for statistical bounds relating the signal strength to the probability of detection).

### C. Loop Counting Asymmetric (LCA)

As mentioned above, we can combine the column-splitting technique above with any BSR-biclustering algorithm in order to search for BAR-biclusters. Here we combine the column-splitting technique with the LCLR algorithm to form what we call the LCA (i.e., Loop Counting Asymmetric) algorithm. We choose the LCLR algorithm for two reasons. Firstly, it has detection-thresholds which are similar to (or better than) many other commonly used biclustering methods, such as Louvain Clustering and UMAP [19], [20] (see Appendix A). Secondly, the LCLR-method naturally provides p-values for each bicluster as a measure of statistical significance (see [18] and [29]).

In contrast to the original LCLR algorithm, where the individual matrix-entries of *B* are drawn from {−1, +1}, in the LCA algorithm the entries of *B* are drawn from {0, +1}. Thus, for each loop of *B*, the score is still organized as the product of all four entries within, but with a different meaning: each nontrivial loop of *B* corresponds to a particular quadrant associated with two distinct columns of *D*. Consequently, each loop of *B* contributes to the score of three of the four different types of BAR associated with those two columns.

We refer to an individual application of the LCLR-method to the split data as algorithm 1.

#### Algorithm 1 Loop Counting Asymmetric (LCA) Row- and Column-Ordering

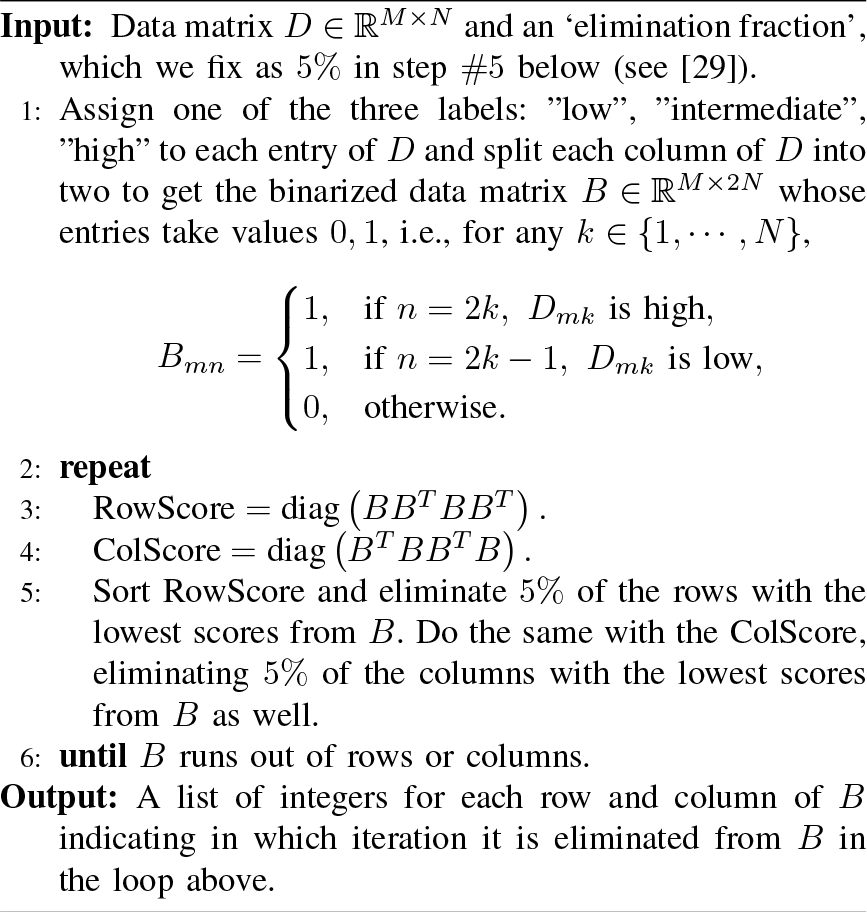

The complete BAR-biclustering algorithm, based on multiple applications of algorithm 1, is presented below in algorithm 2. This second algorithm extracts multiple LCA-biclusters from the data matrix, using the automatic bicluster selection technique described in III-D.

#### Algorithm 2 LCA-Biclustering

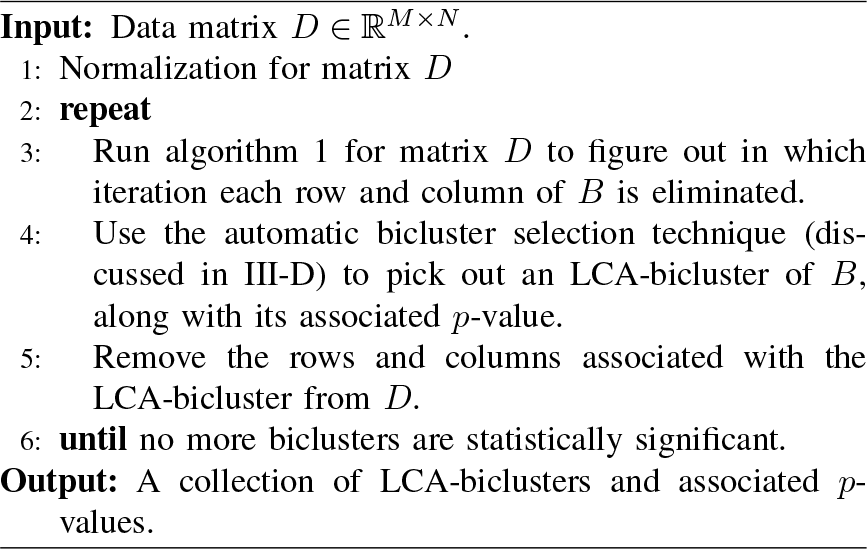

In our numerical experiments below, we’ll use both the LCA- and the original LCLR-algorithm. By comparing these results with one another we’ll be able to determine the degree to which any particular bicluster is BAR- or BSR-specific.

### D. Gene Set Enrichment Analysis (GSEA)

In the numerical experiments, we use both LCA and LCLR to analyze an snRNA-seq data-set from [25]. As we mentioned above, these two biclustering methods search for different kinds of biclusters and reveal different kinds of information within the data-set. To illustrate this point, we collect the gene-subsets of the statistically-significant BAR- and BSR-biclusters that we find, and use gene set enrichment analysis (GSEA) to highlight the statistically-significant gene-pathways that are associated with each bicluster. As we’ll show below, this GSEA returns different subsets of pathways for the different biclusters. Importantly, there are many significant pathways associated with the BAR-biclusters that are not associated with the corresponding BSR-biclusters, indicating that the former contains information not found in the latter.

In terms of implementation, we use the GSEA software from [30]. For each pathway, we compare the enrichment *p*-values for (i) the BAR-biclusters (obtained with the LCA-algorithm) to the enrichment *p*-values of (ii) the trivial set of all genes analyzed, and (iii) the BSR-biclusters obtained with the LCLR-algorithm. We expect to find pathways that have more significant enrichment *p*-values for (i) than for (ii) and (iii).

## III. Data analysis

To illustrate the kinds of BAR-biclusters one might see in practice, and to show that these BAR-biclusters are distinct from the more traditional BSR-biclusters, we apply our LCA-method to the scRNA-seq dataset described in [25]. This dataset comprises 8 different kinds of cell-types drawn from multiple organoids; some of these isogenic-variants have mutations commonly seen in hereditary forms of Alzheimer’s disease. The cells within each cell-type are further divided into 3 stages of development, corresponding to the time that the organoid was allowed to grow before being sequenced.

### A. Fix Cell Type

We apply our biclustering algorithms to each set of samples from each of the cell-types individually. Our goal will be to find (then compare and contrast) the statistically significant BAR- and BSR-biclusters within each cell type. While it is possible for these biclusters to extend across all isogenic variants and all stages of development, we expect at least a few of the biclusters to be biased towards cells (of the fixed type) from a particular isogenic variant or developmental stage. These isogenic- or stage-specific biclusters can be used to highlight genes or gene-pathways that drive heterogeneity within the data-set, and might be linked to Alzheimer-specific genetic mutations and/or development.

### B. Normalize and Binarize

The normalization process mainly consists of truncating the gene-set, based on which binarization is performed. In this particular data-set the vast majority of genes are either not expressed at all, or are rarely expressed across any of the cells in the sample. To focus on those genes that carry the most signals, we limit our analysis to the top 300 genes (approximately the top 1% genes) in terms of overall expression (after eliminating outlier genes with spurious expression values). This choice is made only for the clarity of presentation, and our results do not change qualitatively if we include more genes.

As discussed in [5], the identification of BARs can depend on the choice of thresholds used to define ‘low’, ‘intermediate’ and ‘high’. While more sophisticated studies of BARs can attempt to select different thresholds for different genes, for our proof-of-principle example we use the same thresholding strategy for all genes. Namely, for each gene, we define the lowest 15% of expression-values to be ‘low’, and the highest 15% of expression-values to be ‘high’, with the remainder classified as ‘intermediate’. For normally-distributed data this simple strategy corresponds to defining a threshold proportional to the standard-deviation of the expression-data, as suggested in [5]. Moreover, this simple strategy produces a *B*-matrix which is appropriately centered for subsequent analysis. Once again, our approach is quite robust and the results do not change qualitatively if we choose different thresholds.

### C. LCA & LCLR

We apply both LCA and LCLR to each cell-type array. As described above, the LCA algorithm can reveal biclusters containing a mixture of both BARs and BSRs (the BARs and BSRs can be differentiated at a later stage as described in III-G). By contrast, the LCLR algorithm can only reveal biclusters with a significant BSR-signal. By comparing the results of these two algorithms, we can evaluate how strongly any particular bicluster is driven by BAR-versus BSR-signals. In figure 2, we show an example of the output obtained after running the LCA- and LCLR-algorithms on the data from the Astroglial cells. For this figure we perform a single pass for each algorithm, recording the row- and column-scores in each case. After using the scores to reorganize the rows and columns in descending order, we clearly see significant biclusters in the data (see the block-structure in figure 2C and the low-rank structure in figure 2D, highlighted by the red lines). These correspond, respectively to two large BAR- and BSR-biclusters. In this example we identify altogether four biclusters ‘by hand’, while a more robust and reproducible bicluster selection technique will be described in later contexts. We calculate two dominant principal-components (PCs) of the BAR- and BSR-biclusters, and project each cell’s expression data onto these two PCs. The resulting scatterplots are shown in figure 2E and 2F. These scatterplots are colored by bicluster membership (see legend). Clearly, the subset of cells within the BAR-bicluster is different from the subset of cells within the BSR-bicluster, indicating that the BAR-biclustering method has found a distinct signal within the data.

**Fig. 2.**
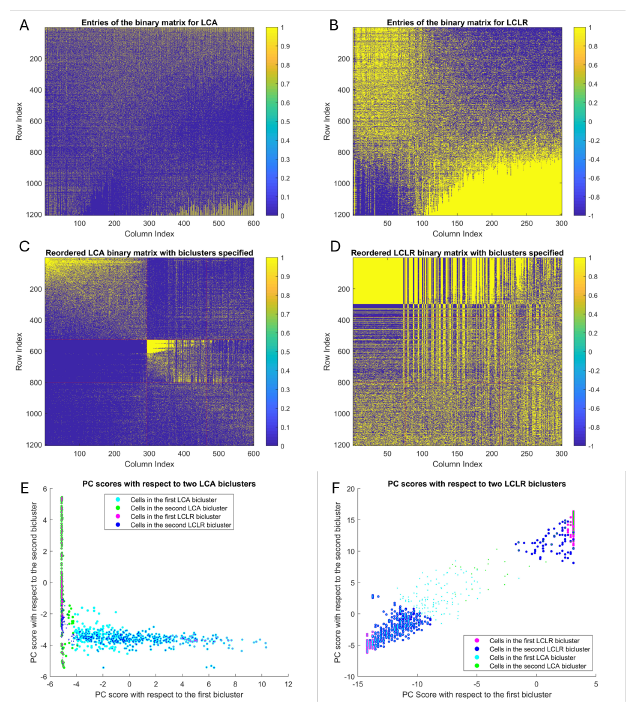
Visualization of LCA and LCLR. (A) the binary matrix for LCA. (B) the binary matrix for LCLR. (C) the reordered binary matrix for LCA. (D) the reordered binary matrix for LCLR. (E) the PC scores with respect to LCA-biclusters. (F) the PC scores with respect to LCLR-biclusters. (A) and (B) show the raw matrices, without any particular structures delineated. (C) and (D) show the reordered matrices, with two biclusters delineated by hand via red lines. (E) illustrates a scatterplot where each cell is projected onto the dominant principal-components of the first two BAR-biclusters found using the LCA algorithm. (F) illustrates a similar scatterplot using the BSR-biclusters found using the LCLR algorithm. In both (E) and (F) the coloring of the dots indicates bicluster membership. Note that the subset of cells delineating the BAR-biclusters is not the same as the subset of cells delineating the BSR-biclusters.

### D. Automatic Bicluster Selection

While the biclusters shown in figure 2 are quite easily delineated by hand, many of the biclusters observed in a typical data-set will be less clear. Consequently, we need an automated method for determining the ‘boundary’ of each bicluster, i.e., which rows and columns define it.

For this purpose, we measure the mean-squared-correlation (MSC) at each iteration of the LCA- and LCLR-algorithms [29].

For any matrix *A* with size *m* × *n*, its MSC is specified as:

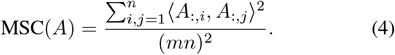

where ⟨ ·, · ⟩ is the standard inner product. Note that the MSC can be re-expressed as:

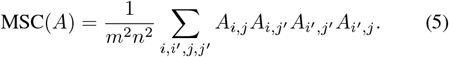

which is proportional to the sum of the row- or column-scores calculated by the LCLR method.

For a matrix *D* with entries in {−1, 0, +1}, the MSC(*D*) is a direct measure of the average squared-correlation between any two rows or columns of *D*; the MSC will be high if the matrix *D* is numerically low-rank. For binary matrices *B* with entries in {0, 1}, the MSC(*B*) will be high if many of the rows or columns of *B* ‘overlap’ (i.e., share many nonzero entries). Because of the way we split the columns of the original data *D* to form *B*, each pair of overlapping columns of *B* corresponds to a particular quadrant associated with two genes of *D*; the MSC of *B* will be high if the matrix *B* is associated with an abundance of BSRs or BARs.

Here we remark that, because the MSC serves as an indicator of both BSRs and BARs, it is natural to measure MSC(*B*) at each iteration of algorithm 1. By comparing this MSC(*B*) with a similar measurement performed on the appropriate subset of the original data-matrix *D*, we should be able to detect if the matrix *B* has an abundance of BARs. We make this intuition more formal below by first introducing some notations.

Define *D* to be the (thresholded) data-matrix. Recall that *D* has entries in {−1, 0, +1}, and LCLR-algorithm is applied on *D*. Recall that the binary matrix *B* used in the LCA-algorithm has entries in {0, 1}, and is obtained by splitting the columns of *D*. We denote this relationship via the ‘column-map’ *µ*:

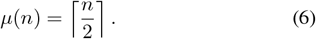

such that any column *n* of *B* corresponds to the gene associated with column *µ*(*n*) of *D*.

When running algorithm 1 on *B*, we iterate multiple times, removing rows and columns within each iteration. Let *l* denote the iteration-index, and *ℳ* (*l*; *B*) denote the row-subset of *B* remaining after iteration *l*. Similarly, let *𝒩* (*l*; *B*) denote the column-subset of *B* remaining after iteration *l*. Finally, let’s refer to *B*(*l*) as the sub-matrix of *B* remaining after iteration *l*. That is, using ‘matlab-notation’:

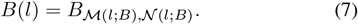

In order to extract the sub-matrix of *D* that corresponds to *B*(*l*) we will use the same row-subset, as well as the corresponding genes, this subset of genes in *D* can be constructed via the column-map: *µ*(*𝒩* (*l*; *B*)). Note that this column-map can (and often does) produce fewer columns than were originally in *𝒩* (*l*; *B*).

Using the row-subset from *ℳ* (*l*; *B*) and the gene-subset from *µ*(*𝒩* (*l*; *B*)), we define:

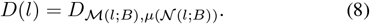

which leads to the definition of MSC-ratio as:

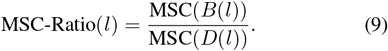

In practice, after running algorithm 1, we measure the MSC-ratio for each iteration *l*.

We assess this empirical MSC-ratio under the null-hypothesis that the data-array is unstructured. To draw a sample from this null-hypothesis, we permute the entries of the normalized data-matrix and rerun the LCA-algorithm on the permuted data. This produces one sample trial from the null-hypothesis, and several such trials are made to estimate the distribution of the MSC-ratio under the null-hypothesis. Once sufficiently many trials have been made, we can use the distribution of MSC-ratios under the null-hypothesis to determine the (empirical) *p*-value of the MSC-ratio of the original data.

Moreover, the permuted trials also allow us to delineate the boundaries of the bicluster within the original data. To be more specific, we first calculate Z-scores for the empirical MSC ratio, with respect to the distribution of MSC ratios drawn from the null-hypothesis (denoted by MSC Ratio Zscore(*l*)). Once we have computed the Z-scores, we define the bicluster using the iteration *l*opt which maximizes this Z-score:

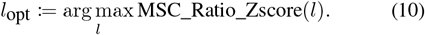

As a result, *B*(*l*_opt_) is the bicluster to select. See figure 3 for an illustration.

**Fig. 3.**
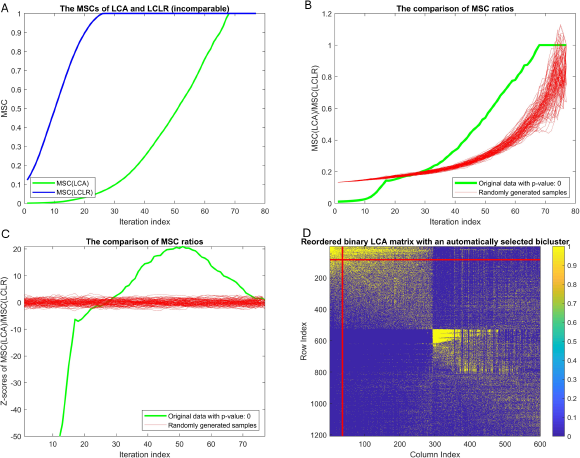
Visualization of automatic bicluster selection. (A) the MSC behavior for LCA and LCLR. (B) the MSC ratio behavior and the randomly generated samples. (C) the Z-scores of the the MSC ratio shown in (B). (D) the automatic bicluster selection result. As (A) illustrates, for the loop-counting algorithm, the MSC always increases and ends up at 1. (B), (C) show curves of the MSC ratio for the given dataset (green) and 100 randomly sampled datasets (red). (D) shows the selected bicluster as the upper left corner of the two red straight lines.

### E. Pearson Test

After delineating the boundaries of the LCA-bicluster, we use the information from the isogenic- and stage-labels (which were not used in the LCA- or LCLR-algorithms) to test whether the bicluster is enriched for any particular isogenicvariant or developmental-stage. The null hypothesis is that the isogenic- and stage-labels are uncorrelated with the LCA-bicluster membership. A Pearson test applied to this hypothesis provides the following results in figure 4. Note that the Pearson test is carried out for the first LCA-bicluster selected consisting of Astroglia cells. The “APP” isogenic-label and the “6-weeks” stage-label are significantly enriched in this LCA-bicluster.

**Fig. 4.**
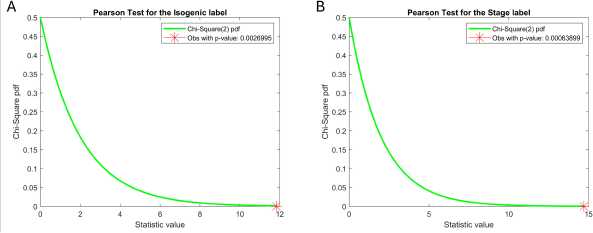
Result of Pearson test. (A) the result of Pearson test on isogenic-labels. (B) the result of Pearson test on stage-labels. The green curve stands for the *χ*^2^ density with degree of freedom 2, while the red star represents the value of the *χ*^2^-statistic computed for the selected LCA-bicluster. In this case, two tests are both statistically significant at the level *α* = 0.01.

In the case above, the distributions of isogenic- and stage-labels within the selected LCA-bicluster are both statistically different from those within the whole data-set, implying that the BAR-bicluster illustrated in this example is indeed specific to certain isogenic-variants and certain developmental-stages. The specific enrichment of this BAR-bicluster is also different from the enrichment of the BSR-biclusters found within the same data-set.

### F. GSEA Results

To further investigate the BAR-biclusters found by the LCA-algorithm, we use GSEA on each of the respective gene-sets. We use the ‘seek’ software from [30], referencing the gene on-tologies `go_bp_iea`, `go_cp_iea`, and `go_mf_iea`, which have each been curated by the seek developers. Our baseline level of significance is calculated using LCLR or by simply inputting all relevant genes we analyzed (the trivial case). We only list pathways that are more significant in the BAR-biclusters but are less or even not significant in any of the BSR-biclusters.

It is typical for the LCA-algorithm to identify gene-subsets that are more strongly enriched than the LCLR-algorithm. One example, for a specific set of biclusters, is shown in table I and table II.

**TABLE I.** Part of GSEA results on BAR-biclusters compared to the trivial case.

| Gene Ontology | Gene Pathway | BAR-bicluster | trivial case |
| --- | --- | --- | --- |
| go_bp_iea | Negative Regulation of Transcription DNA Dependent | $5.9 \times 10^{-5}$ | $1.1 \times 10^{-2}$ |
| | Generation of Neurons | $4.4 \times 10^{-4}$ | - |
| go_cp_iea | Clathrin Sculpted Vesicle | $5.7 \times 10^{-4}$ | $2.7 \times 10^{-3}$ |
| | Neuron Part | $1.9 \times 10^{-5}$ | - |
| go_mf_iea | Structural Constituent of Cytoskeleton | $3.2 \times 10^{-5}$ | $1.1 \times 10^{-4}$ |
| | GTP Dependent Protein Binding | $6.1 \times 10^{-4}$ | - |

For each gene ontology, two pathways are selected to support our conclusions. With the pathway above to be more significant in BAR-biclusters and the pathway below to be only appearing in BAR-biclusters. The last two columns show respective $p$ -values of gene pathways.

**TABLE II.** Part of GSEA results on BAR-biclusters compared to BSR-biclusters.

| Gene Ontology | Gene Pathway | BAR-bicluster | BSR-bicluster |
| --- | --- | --- | --- |
| go_bp_iea | Negative Regulation of RNA Metabolic Process | $8.3 \times 10^{-5}$ | $1.6 \times 10^{-3}$ |
| | Negative Regulation of Transcription DNA Dependent | $5.9 \times 10^{-5}$ | - |
| go_cp_iea | Mitochondrial Envelope | $2.6 \times 10^{-9}$ | $1.8 \times 10^{-6}$ |
| | Nuclear Chromosome Telomeric Region | $3.5 \times 10^{-3}$ | - |
| go_mf_iea | Macromolecule Transmembrane Transporter Activity | $9.3 \times 10^{-4}$ | $1.1 \times 10^{-3}$ |
| | Oxidoreductase Activity | $3.6 \times 10^{-5}$ | - |

For each gene ontology, two pathways are selected to support our conclusions. With the pathway above to be more significant in BAR-biclusters and the pathway below to be only appearing in BAR-biclusters. The last two columns show respective $p$ -values of gene pathways.

### G. Construction of Boolean Asymmetric Relationship Network (BARN)

One of the advantages of BAR-signals and BAR-biclusters is that the individual BARs are easily interpreted. As mentioned above, there are 2 different kinds of BSRs: namely positive and negative correlations. There are also four more kinds of BAR-specific interactions defined in I-A. With these interpretations in mind, we can represent each of the biclusters produced by our LCA-analysis as a gene-gene interaction network. Unlike more traditional network representations (which only illustrate correlations and anti-correlations), our diagrams also highlight the BARs between the genes involved. A simple cartoon of a BARN (indicating the symbols used for each interaction) is shown in figure 5, with subsequent figures 6 and 7 illustrating the BARNs corresponding to two of our BAR-biclusters.

**Fig. 5.**
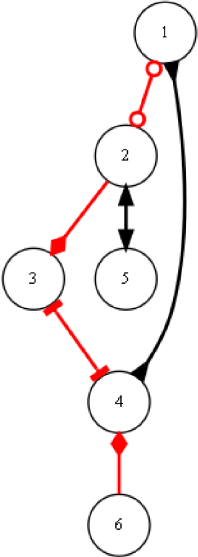
Different types of edges in BARN. The edge between 2 and 5 is a positive correlation, while the edge between 1 and 4 is a negative correlation. Both of them are BSRs, represented by a black edge with the same arrowhead and arrowtail. The edge between 1 and 2 is an ‘nand’ relationship, while the edge between 3 and 4 is an “or” relationship, both of which are undirected BARs. The edge from 2 to 3 represents 2 ⇒ 3, while the edge from 4 to 6 represents 6 ⇒ 4, both of which are directed BARs. BARs are represented as red edges in the network.

**Fig. 6.**
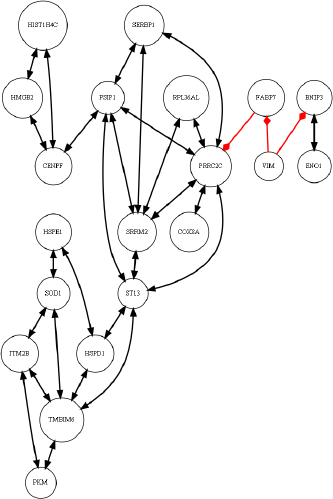
A BARN example. The nodes represent genes, with their names specified. Only the most significant edges are drawn in the network, i.e. those with total-variation-values in the bottom 5%. The edges represent BARs or BSRs as defined above. Red edges are BARs while black edges are BSRs. This BARN is constructed based on the second BAR-bicluster of Glutamatergic neurons.

**Fig. 7.**
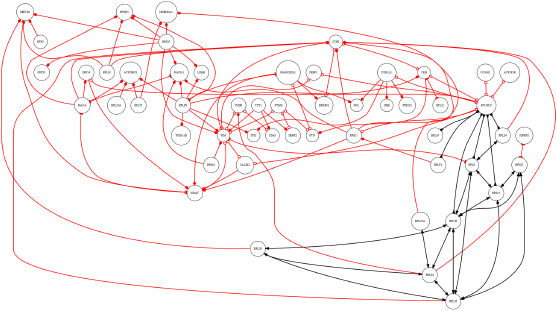
A BARN example with an abundance of BAR. The nodes represent genes, with their names specified. Only the most significant edges are drawn in the network, i.e. those with total-variation-values in the bottom 5%. The edges represent BARs or BSRs as defined above. Red edges are BARs while black edges are BSRs. This BARN is constructed based on the fifth BAR-bicluster of GABAergic neurons.

Note that the LCA-algorithm focuses on biclusters with an overall abundance of BARs and BSRs; the LCA-algorithm by itself will not necessarily identify any particular BAR between any two genes. To determine which particular BARs are over-represented in any of the statistically-significant BAR-biclusters, we apply the following procedure.

First, recall that gene *n* of the original matrix *D* is split into columns 2*n* − 1 and 2*n* within the binary-matrix *B*, corresponding to values of *D*_:,*n*_ being negative and positive, respectively. Given this construction, let’s label the columns 2*n* − 1 and 2*n* (within *B*) as *n*_−_ and *n*_+_, respectively.

Now, given any pair of genes *n* and *n*^*′*^ within *D*, there will be 4 corresponding columns of *B*, denoted by 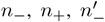 and 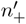 . These four columns of *B* in turn correspond to four nontrivial column-pairs: 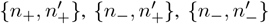 and 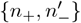 (note that the other two column-pairs {*n*_−_, *n*_+_} and 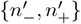 are orthogonal by construction). These nontrivial column-pairs correspond, respectively, to quadrants 1 through 4 of the two-dimensional plane associated with genes *n* and *n*^*′*^.

The inner-product 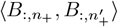 measures the (empirically observed) number of observations of the {*n, n*^*′*^} genepair which fall into quadrant 1. Similarly, the inner-product 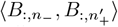 measures the number of observations which fall into quadrant 2, and so forth. Thus, given any gene-pair {*n, n*^*′*^} and any bicluster, we can use the inner-products of the appropriately chosen columns of *B* (within the bicluster) to determine the empirical joint-distribution *ρ*_*n*,*n′*_ (*x, y*) corresponding to the two-dimensional plane associated with that gene-pair (i.e., across values of *x* and *y* ∈ {−, +}).

If we are to observe a joint-distribution with the shape:

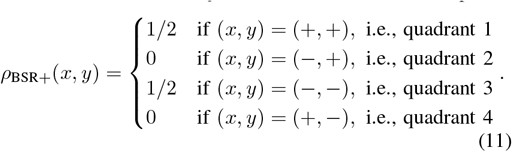

then that joint-distribution would be a perfect BSR, corresponding to ‘positively correlated’ observations within the bicluster. Similarly, if the joint-distribution were to look like:

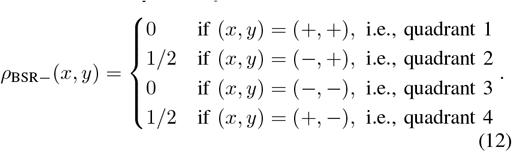

then the gene-pair would be perfectly ‘negatively correlated’ within the bicluster.

Continuing in the same vein, we can define the idealized distribution:

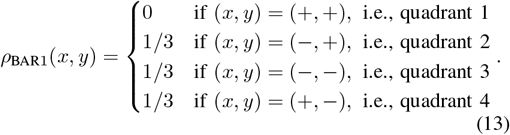

as the BAR for which quadrant-1 is vacant (i.e., the ‘nand’ BAR). We can define the idealized joint-distributions *ρ*_BAR2_ through *ρ*_BAR4_ for the other quadrants analogously.

To assign a relationship to any particular gene-pair {*n, n*^*′*^}, we measure the empirical joint-distribution *ρ*_*n*,*n′*_ as described above, and then compare *ρ*_*n*,*n′*_ to each of the six different idealized joint-distributions *ρ*_BSR+_, …, *ρ*_BAR4_. We use the idealized joint-distribution closest to the empirical joint-distribution to determine the type of BSR or BAR to assign to that gene-pair, using the total-variation as a measure of the difference between distributions. We also record the total-variation associated with each BSR- or BAR-label; by thresholding these total-variation values we can visualize a pruned BARN retaining only the most relevant edges.

These BAR-bicluster-informed networks are similar to many of the boolean-networks described in [15], [31]–[33]. The main difference is that the BARs and BSRs displayed in the network are identified through biclustering, rather than through a direct pairwise analysis of genes across the entire cell-population. As a consequence, gene-gene relationships highlighted in the BARN are significant within only a subset of the cells examined, and are typically *not* expressed across the entire cell-population.

We point out that, while it is certainly possible to have a network comprised only of BSRs (e.g., a collection of genes that are all driven by a single upstream source), it is typically not the case that BAR-biclusters are comprised only of BARs. This is because a large interconnected network of genes with multiple BARs will typically give rise to emergent correlations between certain genes in the network. For example, if gene-A and gene-B are in an ‘or’-type BAR, and gene-B and gene-C are also in an ‘or’-type BAR, then genes A, B and C could form a BAR-bicluster. This BAR-bicluster typically includes the BSR-relationship (a positive correlation) between genes A and C.

### H. A Prediction Use Case

As described above, the BAR- and BSR-biclusters contain different subsets of cells and highlight different kinds of relationships. An immediate question is whether or not the additional information afforded by the BAR-biclusters is useful. To illustrate just one potential application of BAR-biclustering, we demonstrate that the BAR-signals discovered by the LCA-algorithm can augment the standard LCLR-algorithm and improve classification. As a first step, we randomly divide the original data-set into two groups of cells (i.e., by row). We will use one of these groups as a ‘training’ group for the LCA- and LCLR-algorithms. After identifying biclusters within the training group, we use the BSR-signals within these biclusters to build a BSR-only linear-classifier for the isogenic- and stage-labels within the training group. We also use the BSR- and BAR-signals together to build a BAR + BSR linear-classifier for the training group. To simplify the model, we only consider a binary classification problem, e.g., fix a stage-label denoted as “Stage 1”, any prediction would be “is Stage 1” or “is not Stage 1”. Finally, we validate these two linear classifiers within the second ‘testing’ group.

In terms of details, we build a linear-classifier by first measuring the dominant principal-component (PC) of each statistically-significant BAR- and BSR-bicluster within the training set. Then, each cell is projected onto these PCs to be assigned a bicluster-specific score. We use logistic-regression to link these bicluster-specific scores to the ground-truth isogenic- or stage-labels (within the training data). Finally, we use the output of this logistic-regression (i.e., the logistic link function) as a predictor of label within the testing data.

As expected, the linear-classifier we build using information from both the BAR- and BSR-biclusters outperforms the classifier built using only the BSR-biclusters. Shown in figure 8 is the ROC-curve associated with each classifier. While this is only a single example, we see similar results throughout our analysis. The same general trend holds: BAR-signals provide extra information which can be used to augment BSR-signals and build better classifiers. We remark that our strategy is only one of many, and other methods may be even more effective at predicting isogenic- or stage-label within this data-set.

**Fig. 8.**
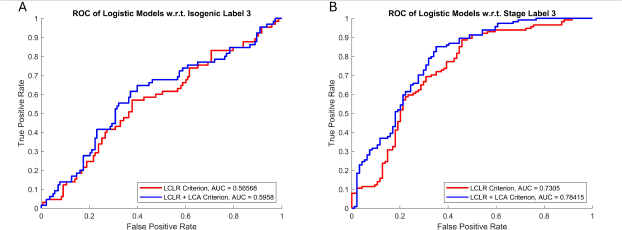
BAR as a supplement of BSR in prediction. (A) prediction on whether a cell has the third isogenic-label. (B) prediction on whether a cell has the third stage-label. After adding BAR information, the AUC is higher. The performance of the predictions for the other isogenic- and stage-labels are qualitatively similar.

## IV. Conclusion and discussion

We have introduced a very simple but effective strategy for finding BAR-biclusters. The essence of this strategy involves splitting each column of the original data-set into two, allowing for individual quadrant information to be accessed using standard inner product calculations. This simple preprocessing step can be followed by any biclustering method that searches for BSRs. Above we demonstrated one simple BAR-biclustering method: – the LCA-algorithm – built using the LCLR-algorithm of [18]. While we believe that the LCLR method is a good choice for biclustering general data-sets, we remark that an advantage may be gained by choosing a biclustering method which is appropriate for the domain of application and the specifics of the data-set at hand.

We use our LCA-algorithm to analyze an scRNA-seq dataset from [25]. As described above, the LCA-algorithm is capable of finding multiple isogenic- and stage-specific BAR-biclusters. Importantly, the cell- and gene-membership of these BAR-biclusters are quite different from the membership of the corresponding BSR-biclusters (found using the standard LCLR-algorithm). The BAR-biclusters can be used to construct BAR-networks, which illustrate new and potentially useful groups of gene-gene interactions. Furthermore, the BAR-signals revealed by the LCA-algorithm can be used to build better linear-classifiers for predicting the isogenic- and stage-labels; these improved classifiers might be useful for better categorizing future data-sets.

Finally, we mention that our column-splitting strategy can be modified to include a variable-specific weight to each column. For example, after centering the data and splitting into positive- and negative-parts, one might retain the absolute-value of each part, rather than encoding the positive- and negative-parts with the value of 1 [55]. This coding strategy could allow for larger data-values to play a more dominant role in the subsequent biclustering step. We leave the analysis of this method to future work.

### A. Back to the Organoid Dataset

Last but not least, we take a final look at the GSEA results for the BAR- and BSR-biclusters found within the snRNA-seq dataset of [25]. As commented on previously, most of these biclusters are enriched for some of the isogenic-variants. We can label the biclusters enriched for the APP- and PS1-variants as ‘AD-specific’, and the biclusters enriched for the wild-type as ‘non-specific’ (or ‘WT-specific’). With these labels in place, we can look for gene-pathways which are more strongly enriched within the AD-specific BAR-biclusters than the non-specific biclusters. Some results of this search are shown in table III, with the remaining results shown in Appendix B. These results are suggestive, as many of these pathways may play a role in the differences between AD- and WT-organoids seen in [25].

**TABLE III.**
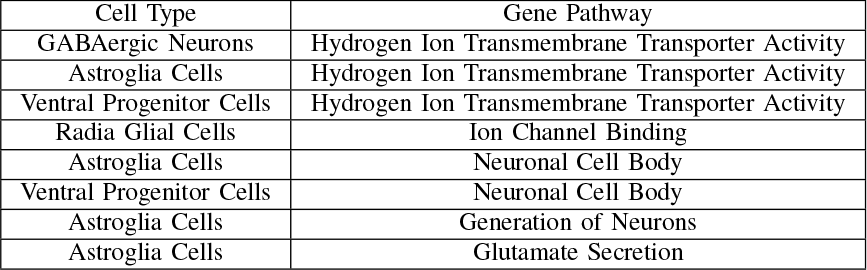
Difference in significant gene pathways from WT- and AD-biclusters.

For example, the genes and pathways related to ion-transport and ion-channel binding may be partially responsible for the differences in conductivity and current-density observed between the isogenic-variants. Similarly, the pathways related to neuronal cell generation might play a role in the differences in arborization observed between the AD- and WT-organoids. As another example, the most significant BARs (red-arrows) in figure 6 highlight genes such as vimentin and FABP7; the former encodes for filament proteins responsible for cytoskeletal structure and function, while the latter is a fatty-acid binding protein relevant for glial-fiber development in the brain. Both have been associated with AD in several previous studies [45]–[52].

## V. Acknowledgments

This research was supported by the following NIH grants: 3U19AG023122-13S2, UH3AG064706 and U19 AG065169.

## Appendix A

### More details on LCLR

Case studies on the comparison between LCLR, UMAP and Louvain clustering.

## Appendix B

## Notes

https://github.com/Haosheng-Zhou/BAR-LCA

### Competing Interest Statement

The authors have declared no competing interest.

### Summary of Updates

Author updated; Introduction updated to cite more recent works.

