## Appendix A for "Detecting Boolean Asymmetric Relationships with a Loop Counting Technique and its Implications for Analyzing Heterogeneity within Gene Expression Datasets"

### More Details on LCLR

#### A.1 Comparing the LCLR method against Umap and Louvain clustering:

In the main text we reference the 'LCLR' biclustering method, also referred to as the 'loop-counting' method. In this appendix we compare this loop-counting method (as described in [5]) to two other commonly used clustering algorithms: umap-clustering and louvain-clustering (implemented in [10, 1] and [2], respectively).

First, we use simulated data to demonstrate that the loop-counting method can be more sensitive than umap- and louvain-clustering. Second, we demonstrate that the loop-counting method performs acceptably on a real scRNA-seq data-set from [4].

##### A.1.1 Implementation details:

The main difference between the loop-counting algorithm we use here and the one described in the main text is that, in this appendix, we will only implement the binary-symmetric-biclustering methods described in [5]. In addition, we will repeatedly use this loop-counting algorithm to subdivide the data-set into two groups, forming a hierarchical tree where each branch corresponds to a group of cell samples.

In more detail: At each branch of the tree (starting with the entire data-set) we use the strategy of [5] to order the remaining samples based on their average sample-sample correlations with the other samples in that branch. This process is almost identical to the automatic cluster selection described in the main text. The only difference is that, because we are only searching for BSR-biclusters, we use the the average sample-sample correlations from the LCLR iterations directly, rather than calculating the MSC-ratio described in the main text. Additionally, each time we order the samples of any branch of the tree, we also estimate the statistical-significance of the observed sample-sample correlations (referred to as the 'trace' in [5]).

Our null-hypothesis 'H0' is that the samples are drawn from an anisotropic-gaussian-distribution with the empirically-observed transcript-transcript correlations, and we can easily draw from this null-hypothesis by randomly-rotating the remaining samples within sample-space. By comparing the original samples to the distribution of samples drawn under the null-hypothesis, we extract two important pieces of information. First, we estimate the p-value for the current branch – i.e., the probability $p$ that the observed correlations could have originated from samples that were drawn from H0. Second, we use the observed trace to split the current set of samples into two groups, choosing the split which is least likely to have originated from the null-hypothesis. By construction, at least one of these groups will have an average correlation which is statistically significant with respect to H0 (i.e., with a p-value at most $p$).

Thus, ultimately, our p-value for any given branch at any point along the tree is an estimate of the probability of observing the sequence of splits producing that branch, conditioned on the assumption that – once any split is performed – the data on each side is gaussian-distributed with a gene-gene covariance-matrix which matches that observed empirically. This is certainly not the same thing as the *unconditioned* probability of observing the sequence of splits producing that branch from a gaussian starting distribution, but we believe that it is a reasonable approximation in many real-world situations. One piece of corroborating evidence is that, when testing our algorithm, we rarely see multiple spurious recovered-clusters (see Fig A.5 below), implying that our p-value estimate is not too far from the unconditioned probability mentioned above.

We then proceed recursively, repeatedly subdividing each branch of the tree until no further statistically-significant subdivisions can be made. The p-value of the branches increases along any path in the tree, and the entire tree can be arranged so that each level corresponds to a single p-value. Consequently, once the tree is constructed, a user can fix a p-value-threshold (e.g., 0.05) and read off the clusters associated with that threshold – each of these clusters will in turn correspond to a path along the tree which is at least as significant as the chosen threshold. Furthermore, any clusters which persist over a wide range of p-values will be robust (in the sense of our null-hypothesis H0 above).

##### A.1.2 Simple Case Study: random-matrix with planted-clusters

As an example of our approach, we'll describe a very simple case-study involving a random-matrix with several 'planted-clusters'. To start with, we create a random matrix $A$ of size $M$ by $N$, with the rows and columns representing samples and genes, respectively. Each entry of $A$ will be drawn independently from a standard-normal-distribution. After creating $A$, we'll fix a 'signal-to-noise' level $\sigma$, and adjust the entries of $A$ to implant $K$ different clusters into $A$. Each cluster will have a signal-to-noise ratio of $\sigma$, and will be associated with a different set of samples and genes. To implant cluster-$k$ of size $m[k] \times n[k]$ into $A$, we define the cluster-mean $\mu[k] = \mu(m[k], n[k]) = \sigma \cdot \sqrt{N}/\sqrt{m[k]n[k]}$, and then increase the entries of each column of $A$ corresponding to that cluster by $\pm\mu[k]$ (with the sign chosen at random for each column). This ensures that the genes associated with any particular cluster are differentially-expressed with respect to the remainder of $A$. We use the term 'signal-to-noise' to refer to $\sigma$ because it is the ratio between the dominant eigenvalue of the 'signal' $\mu[k]$ associated with any individual cluster and the dominant eigenvalue of the noise.

We show an example of such a random-matrix in Fig A.1, with $M = 563$, $N = 6325$, $K = 6$, and $\sigma = 0.70$. For this particular example the $K = 6$ planted-clusters are of varying size, containing $m[k] = 7$, 12, 19, 31, 51 and 85 samples, respectively, with the remaining 358 samples forming a 'background'-cluster. The number of genes in each planted-cluster is $n[k] = 14$, 28, 57, 113, 228, 458 and 458, respectively, with the remaining 5427 genes left to the background-cluster. Due to this particular construction, we know the ground-truth: there are $K + 1 = 7$ different clusters within the data: the $K = 6$ planted-clusters described above, as well as the 'background' cluster consisting of the remaining samples.

The typical steps of an scRNA-seq analysis involve first projecting the data-matrix down to a handful of dimensions, and then applying an unsupervised clustering-algorithm to the projected-data. Thus, for this particular example we'll consider projecting onto the $R = 7 + 2 = 9$ dominant principal-components, which should be sufficient to capture the $K + 1 = 7$ clusters in the data.

When we apply the hierarchical loop-counting algorithm to this projected-data, we obtain the tree shown on the left of Fig A.2. For this example our algorithm first identifies a split with a very high significance-level

| 563 | 322 | 106 | 67 | 30 | 19 | 12 | 7 |
|---|---|---|---|---|---|---|---|
| 358 | 308 | 33 | 17 | 0 | 0 | 0 | 0 |
| 7 | 0 | 0 | 0 | 0 | 0 | 0 | 7 |
| 12 | 0 | 0 | 0 | 0 | 0 | 12 | 0 |
| 19 | 0 | 0 | 0 | 0 | 19 | 0 | 0 |
| 31 | 1 | 0 | 0 | 30 | 0 | 0 | 0 |
| 51 | 1 | 0 | 50 | 0 | 0 | 0 | 0 |
| 85 | 12 | 73 | 0 | 0 | 0 | 0 | 0 |

Table A.1: This is the confusion-matrix representing the intersections between the ground-truth-clusters and our recovered-clusters for the loop-counting algorithm applied to the example shown in Fig A.1 after projecting onto $R = 9$ principal-components. The total number of samples is $M = 563$, shown in the upper left corner. The rows of the table correspond to the ground-truth-clusters, whereas the columns correspond to the recovered-clusters. The sizes of each ground-truth-cluster are listed in the first column, while the sizes of each recovered-cluster are listed in the first row. The remainder of the array lists the sizes of the intersections between each of the ground-truth- and recovered-clusters. The collection of recovered clusters has a quality $\log(1/P) \sim 394$. For this example an exact recovery would have a quality $\log(1/P) \sim 550$.

(i.e., $\log(1/p) \sim 27$), followed by 5 more splits of decreasing significance, ending with a split at significance-level $\log(1/p) \sim 7.5$. Our algorithm is set to terminate at $p = 0.05$ by default (corresponding to $\log(1/p) \sim 3$), and so we observe no further splitting. If we had allowed our algorithm to continue all the way to $p = 1$ (corresponding to $\log(1/p) \sim 0$), we would of course continue splitting each group of samples further and further until we had $M$ singleton-groups remaining.

If we were to set $p = 0.05$ as a threshold, we would extract 7 clusters from that level of the tree (i.e., the $\log(1/p) = 3$ level of Fig A.2). These clusters strongly overlap with the 7 ground-truth clusters in the data-matrix, as shown in the 'confusion-matrix' displayed in table A.1. To evaluate the quality of this recovery, we calculate the P-value described in section A.1.5. This P-value measures the probability of obtaining a better result by randomly permuting the recovered sample-labels. For this particular result the P-value is quite small, with $\log(1/P) \approx 394$. This high quality indicates that the recovered-clusters are informative of the ground-truth. The best possible P-value (assuming exact recovery) for this example is $\log(1/P) \approx 550$.

In general, the organization of the recovered clusters depends on which level of the tree we consider. For example, had we set a more restrictive p-value threshold of $\log(1/p) \sim 20$, then we would have only read out 5 recovered-clusters, and the associated quality would have been $\log(1/P) \sim 200$. The relationship between the p-value threshold and the quality $\log(1/P)$ is shown on the right of Fig A.2 in gray.

For this particular example louvain- and umap-clustering (with default parameters) produce 6 and 1 recovered clusters, respectively, with confusion-matrices displayed in tables A.2 and A.3. These recovered-clusters are not as close to the ground-truth as the recovered-clusters shown in table A.1, and have a lower quality, indicated on the right of Fig A.2 in dashed-red (louvain) and dashed-black (umap). The number of recovered-clusters for louvian- and umap-clustering (in this case 6 and 1, respectively) is indicated via the red- and black-inset-text on the right of Fig A.2.

##### A.1.3 Simple Case Study: some commonly observed phenomena

In this subsection we modify the case-study above to investigate the behavior of the various clustering-algorithms. Because our case-study is so idealized, we only vary a few rather general parameters: (i) the signal-to-noise ratio $\sigma$ (related to how easy it is to recover the ground-truth) and (ii) the projection-dimension $R$ (a common parameter used in many analysis-pipelines). As we'll see, the performance of louvain- and umap-clustering can change quite

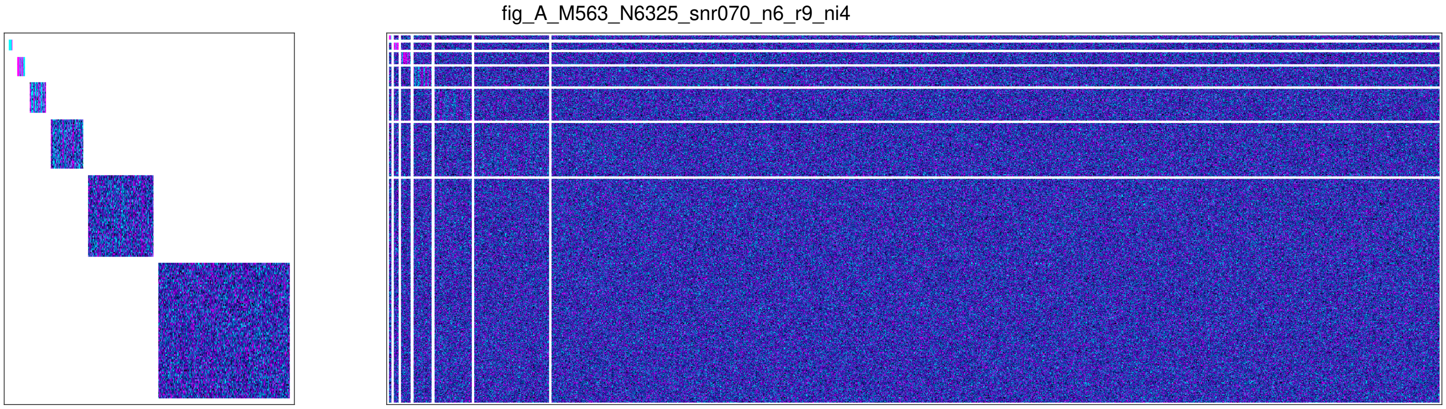


Figure A.1: Here we illustrate a random-matrix with $K = 6$ planted clusters. The clusters are of varying size, each corresponding to a distinct subset of rows and columns, as shown on the left. On the right we show the full matrix $A$, produced by combining the clusters with a random noisy background. In both pictures we have arranged the rows and columns so that the individual clusters are contiguous submatrices of $A$. We have also left gaps (white) between the row- and column-subsets associated with each of the clusters.

| 563 | 164 | 150 | 142 | 57 | 38 | 12 |
|---|---|---|---|---|---|---|
| 358 | 100 | 107 | 94 | 26 | 26 | 5 |
| 7 | 0 | 0 | 0 | 0 | 0 | 7 |
| 12 | 0 | 0 | 0 | 0 | 12 | 0 |
| 19 | 0 | 19 | 0 | 0 | 0 | 0 |
| 31 | 0 | 0 | 1 | 30 | 0 | 0 |
| 51 | 3 | 7 | 41 | 0 | 0 | 0 |
| 85 | 61 | 17 | 6 | 1 | 0 | 0 |

Table A.2: This is the confusion-matrix for the louvain-clustering-algorithm applied to the example shown in Fig A.1 after projecting onto $R = 9$ principal-components. The collection of recovered clusters has a quality $\log(1/P) \sim 152$.

| 563 | 563 |
|---|---|
| 358 | 358 |
| 7 | 7 |
| 12 | 12 |
| 19 | 19 |
| 31 | 31 |
| 51 | 51 |
| 85 | 85 |

Table A.3: This is the confusion-matrix for the umap-algorithm applied to the example shown in Fig A.1 after projecting onto $R = 9$ principal-components. The collection of recovered clusters has a quality $\log(1/P) = 0$ (i.e., no information).

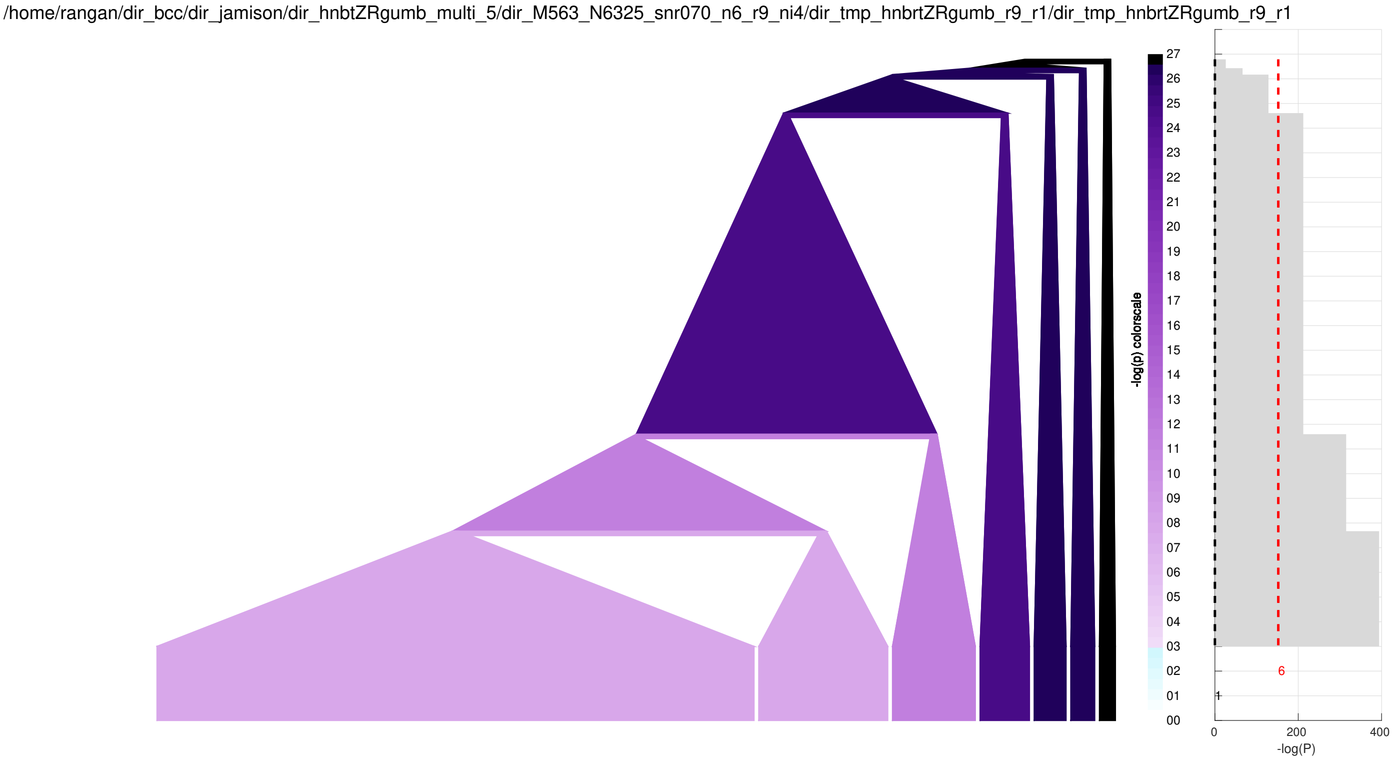


Figure A.2: Here we illustrate the results of the hierarchical loop-counting algorithm applied to the example shown in Fig A.1 after projecting onto $R = 9$ principal-components. On the left we show the tree associated with the hierarchy of clusters. This tree spans p-values ranging from 0 to 0.05 (i.e., $\log(1/p) \geq 3$). The samples are organized along the horizontal axis, with the branches of the tree indicated with triangles. Each branch touching the bottom of the figure contains the samples spanned by its base, and those branches further up the tree contain all the samples of their children (i.e., branches below them). The branches are drawn so that the root of each branch (i.e., the upper point of its triangle) is placed at a vertical-value corresponding to the $\log(1/p)$ of that branch. Additionally, each branch is color-coded based on its $\log(1/p)$-value, as shown via the colormap in the middle. The tree is arranged so that each vertical level (i.e., each horizontal line) corresponds to a single p-value threshold. On the right we show the quality $\log(1/P)$ of the clustering associated with each p-value threshold (gray). We also show the quality of louvain-clustering (dashed-red) and umap-clustering (dashed-black) applied to the same example. The number of recovered-clusters for louvain- and umap-clustering is shown below the dashed-lines in red and black, respectively.

a bit as these parameters are varied. In certain regimes these clustering-algorithms can return multiple spurious clusters, making it difficult to determine which recovered-clusters (or recovered-cluster-combinations) are statistically-significant.

First we vary the signal-to-noise-ratio $\sigma$ within the range $\sigma \in [0.5, 1.0]$, while fixing the projection-dimension to be $R = 9$. For each $\sigma$, we repeat our numerical experiment from the previous section multiple times (over 32 trials). We run the hierarchical loop-counting algorithm, as well as louvain- and umap-clustering on each trial, measuring the number of recovered-clusters, as well as the recovered-cluster-quality $\log(1/P)$. The trial-averaged results are shown in Fig A.3.

Recall that, for this example, there are $K + 1 = 7$ ground-truth clusters. For this particular choice of $R$, louvain-clustering typically returns fewer than 7 clusters, even when $\sigma$ is high. The behavior of umap-clustering is more complicated, returning very few clusters when $\sigma$ is small, and returning the correct number of clusters (i.e., $\sim 7$) when $\sigma$ is very large, but returning a large number of spurious clusters for intermediate values of $\sigma$.

To investigate this phenomenon more carefully, we fix the signal-to-noise-ratio $\sigma = 0.80$ at a point where umap-clustering returns multiple spurious clusters, and vary the projection-dimension $R$. We consider values of $R-(1+K)$ ranging logarithmically from 1 to 512 (i.e., the number of 'excess-dimensions' is 1, 2, 4, 8, etc.). For each choice of $R$ we run 32 trials. The trial-averaged results are shown in Fig A.4.

Note that, when $R$ is small, umap-clustering typically returns a great many clusters that – while difficult to sift through – are still somewhat high in terms of quality. However, as $R$ increases, the number of recovered-clusters returned by umap decreases dramatically, as does the overal quality of umap's results. By contrast, louvain-clustering produces results with a quality that is somewhat insensitive to $R$, but the number of recovered-clusters grows dramatically as $R$ increases.

While a comprehensive study of louvain- and umap-clustering is outside the scope of this appendix, we have observed that the behavior of louvain- and umap-clustering can be quite complicated. In general, their results depend on both $\sigma$ and $R$, as well as the details of the data-set (e.g., the number and size of the ground-truth clusters, how well separated they are from one another, how they are distributed, etc.). We conclude this particular case-study by simultaneously varying both $\sigma$ and $R$ (within their respective ranges mentioned above). The trial-averaged results are shown in Fig A.5.

Note that umap-clustering returns the correct number of high-quality recovered-clusters only when the signal-to-noise-ratio $\sigma$ is quite high and the number of excess-projection-dimensions $R - (1 + K)$ is quite low. As $\sigma$ decreases and $R$ increases the number of umap's recovered-clusters increases dramatically, and the quality decreases. By contrast, louvain-clustering returns recovered-clusters of moderate quality for a wide range of projection-dimensions $R$, provided that the signal-to-noise-ratio is not too low. When $R$ becomes large louvain-clustering tends to return a very large number of recovered-clusters (which are devoid of any information when the signal-to-noise-ratio is low).

In summary, unsupervised clustering-algorithms can sometimes produce results which are confusing; the number of recovered-clusters can differ greatly from the true number of ground-truth clusters, and cluster-quantity is not always an indicator of cluster-quality. We expect that the loop-counting method can be of some use in situations like these. As described above, the loop-counting method can be structured to produce a tree where each branch is associated with a p-value. This tree structure can be helpful in determining which of the recovered-clusters might be important. Specifically, branches that are highly significant will correspond to clear (linearly-discriminable) signals, while those branches that are not particularly significant will correspond to more subtle heterogeneous structures that not that different from an anisotropic gaussian-distribution in sample-space. Along the same vein, branches which persist across a wide range of p-values will correspond to clusters that are robust, while branches that only exist for a limited range of p-values will correspond to clusters that might be difficult to define precisely.

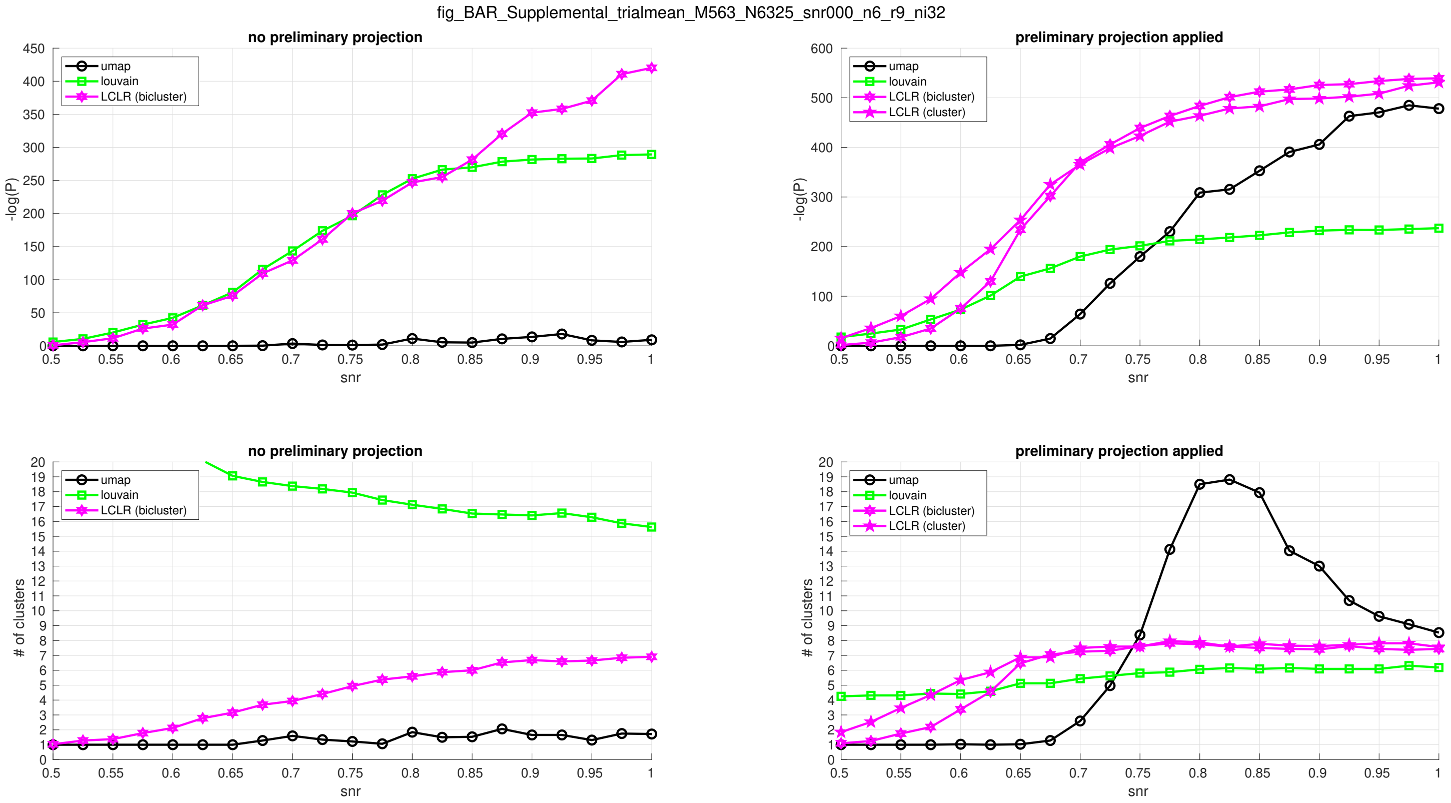


Figure A.3: Here we show the recovered-cluster-quality $\log(1/P)$ (top subplot) and the number of recovered-clusters (bottom subplot) as a function of the signal-to-noise-ratio for the case-study described in section A.1.2. On the left-hand side we show the results with no preliminary-projection. In this context one would run the standard LCLR algorithm, as described above and in [26], and labeled as 'LCLR (bicluster)'. On the right we show results when the projected-rank $R = 9$. In this scenario one might also consider running a modified version of the LCLR algorithm where only samples are removed (and all genes are retained). This modified algorithm is more similar to a standard clustering algorithm, and is labeled as 'LCLR (cluster)'.

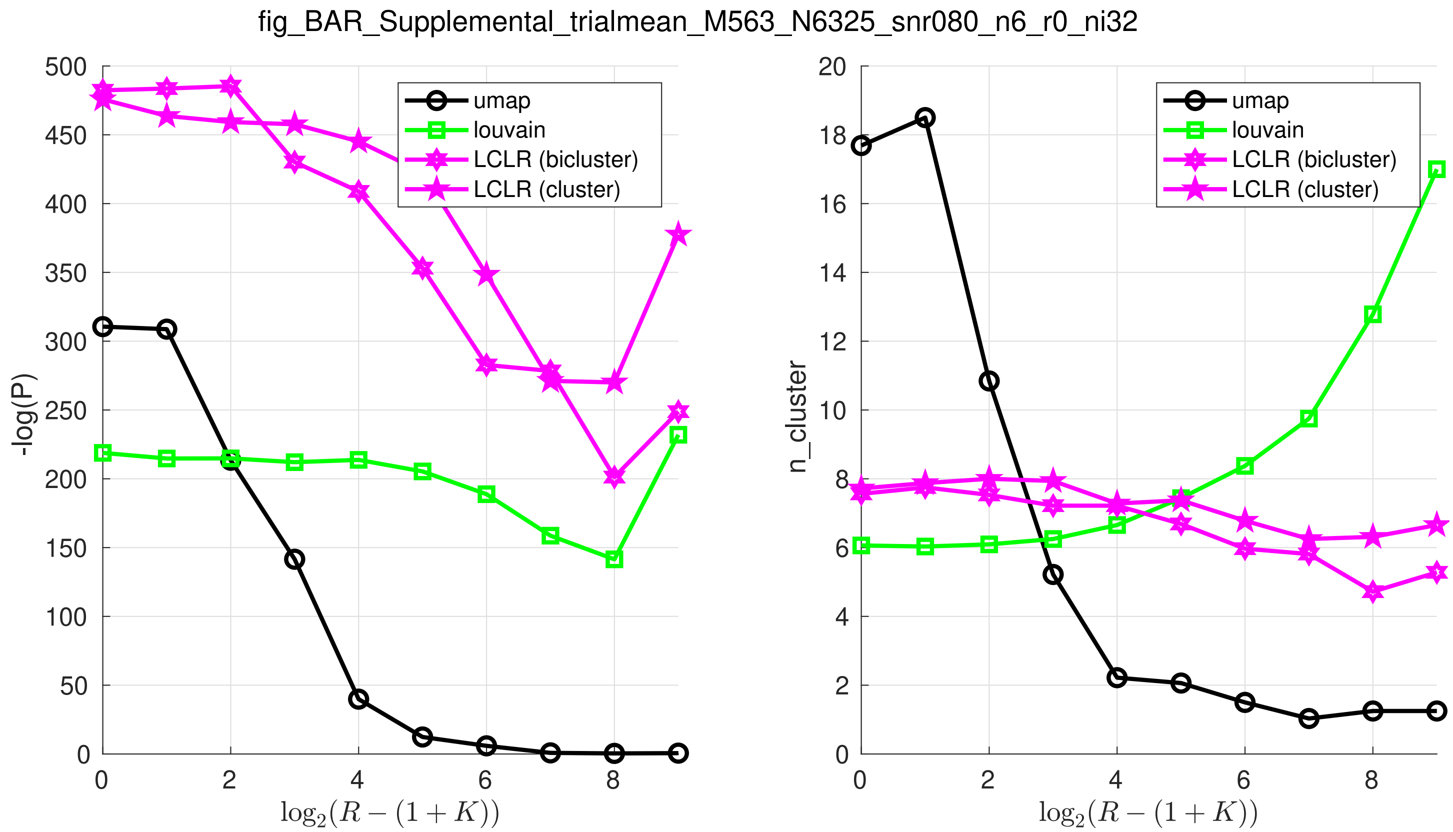


Figure A.4: Here we show the recovered-cluster-quality $\log(1/P)$ (left subplot) and the number of recovered-clusters (right subplot) as a function of the projected-rank $R$ for the case-study described in section A.1.2. In each case the signal-to-noise-ratio $\sigma = 0.80$. As in the previous figure, the 'LCLR (bicluster)' and 'LCLR (cluster)' methods refer, respectively, to (i) the standard LCLR algorithm described in the main text and [26], and (ii) a modified version of the LCLR algorithm where only samples are removed, and all genes are retained.

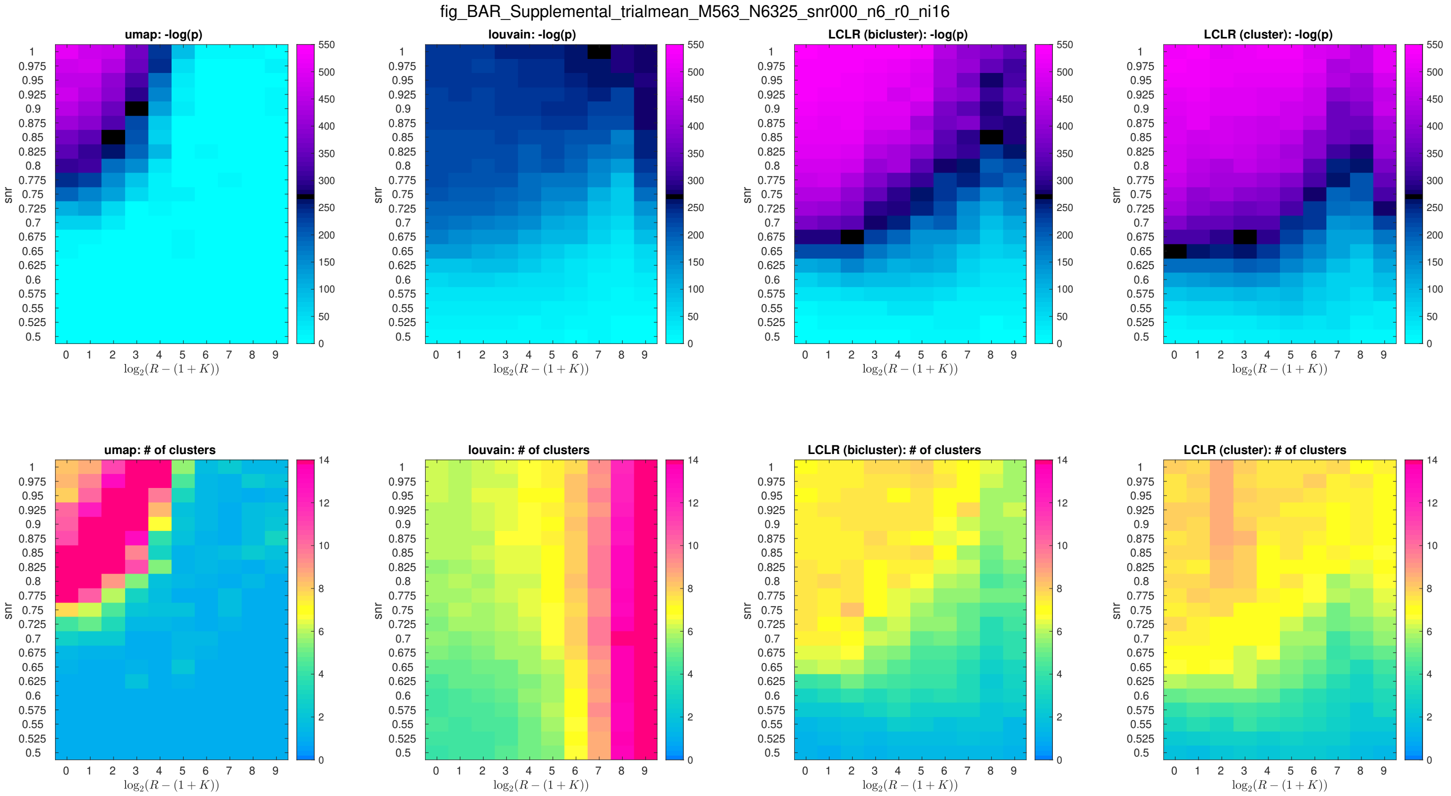


Figure A.5: Here we show heatmaps illustrating the recovered-cluster-quality $\log(1/P)$ (top subplots) and the number of recovered-clusters (bottom subplots) as a function of $\sigma$ and $R$. Each heatmap uses the adjacent colorbar. Recall that for this case-study the number of ground-truth clusters is $K+1=7$ (shown as yellow in the heatmap), and the maximum possible quality is $\log(1/P)\sim 550$ (shown as magenta in the heatmap). The different subplots represent trial-averaged results for the hierarchical loop-counting method, as well as louvain- and umap-clustering.

##### A.1.4 Application to scRNA-seq data:

In this section we demonstrate the performance of the loop-counting method on a single-nuclei RNA-seq data-set involving brain-tissue samples, with the details described in [3, 4]. There are $M = 1781$ samples in this data-set, spanning $N_E = 15137$ exonic-umis, and $N_I = 13760$ intronic-umis. The exonic-data has $R_E = 18$ statistically-significant principal-components, while the intronic-data has $R_I = 20$ statistically-significant principal-components.

We use the published sample-labels from [3, 4] as our 'ground-truth', which defines 46 clusters ranging in size from 194 to 3. These previously published clusters were determined using a supervised pipeline with multiple stages, including several passes of various clustering algorithms such as t-sne [8] which adopts a strategy similar to that of umap [9]. Consequently, we expect that umap-clustering will perform well for this data. However, we remark that, of these ground-truth-clusters, only 17 have a size larger than $\sqrt{M} \sim 42$. This is the detection-threshold of most spectral clustering-schemes applied to homogenous data with a single planted-clique [7], and so we shouldn't expect to recover clusters smaller than this unless they really stand out from the remainder of the data.

While a full scRNA-seq analysis pipeline can involve multiple preprocessing stages including data-normalization, imputation and projection, our goal is simply to compare our hierarchical loop-counting method with umap- and louvain-clustering. Therefore, we adopt the following simple preprocessing stages: We first log-normalize the count-data (after adding a pseudo-count of 1), and we do not impute any of the zero-entries. Subsequently, we project onto the dominant $R$ statistically-significant principal-components, where $R$ is determined via a permutation-test with p-value-threshold of 0.05.

The results for the exons are shown in Figs A.6 and A.7. Note that our hierarchical loop-counting algorithm produces results that are at least as informative as louvain- and umap-clustering applied to the same data-set. Moroever, the tree produced by our method exhibits two wide 'plateaus'; the first ranging from $\log(1/p) \in [15, 25]$, and the second from $\log(1/p) \in [10, 14]$. Across these plateaus the number of recovered-clusters remains relatively constant (i.e., 7 and 13, respectively). These plateaus represent natural cut-points where a user (who does not know the ground-truth) might choose to set their p-value-threshold. The resulting recovered-clusters will be robust in the sense described in section A.1.1: variations of the p-value-threshold (across the span of the plateau) will not greatly perturb the cluster-configuration. Moroever, the branches responsible for the recovered-clusters will each correspond to linear-separations in the data which are at least as significant as the p-value-threshold used.

Results summarizing the introns are shown in Figs A.8 and A.9. Note that, for both the exonic- and intronic-data louvain-clustering produces very few recovered-clusters (i.e., 3 and 4, respectively), while umap-clustering produces many (i.e., 41 and 40, respectively). While umap's recovered-clusters for the exon-data are of reasonably high quality, umap's recovered-clusters for the intron-data are essentially uncorrelated from the ground-truth.

##### A.1.5 Robustness: cluster-comparison

After clustering the samples within any data-set, we often need to compare the results of that sample-clustering with those of previous sample-clusterings, or with a given set of sample-cluster-labels (such as 'ground-truth' labels). We use the simple P-value described below.

To set the stage, assume that we have applied two different clustering-algorithms to the samples. Let's refer to the output of the first clustering-algorithm as the set $\vec{\mathcal{A}}$. The set $\vec{\mathcal{A}}$ is a list

$$\vec{\mathcal{A}} = \{\mathcal{A}[1], \mathcal{A}[2], \ldots, \mathcal{A}[j], \ldots, \mathcal{A}[J]\}$$

of disjoint sets $\mathcal{A}[j]$ which, together, include all $M$ samples (i.e., $\mathcal{A}$ is a disjoint covering of the samples). We'll refer to the number of elements in each set $\mathcal{A}[j]$ as the number $A[j]$, which together form the vector $\vec{A}$. Similarly, let's

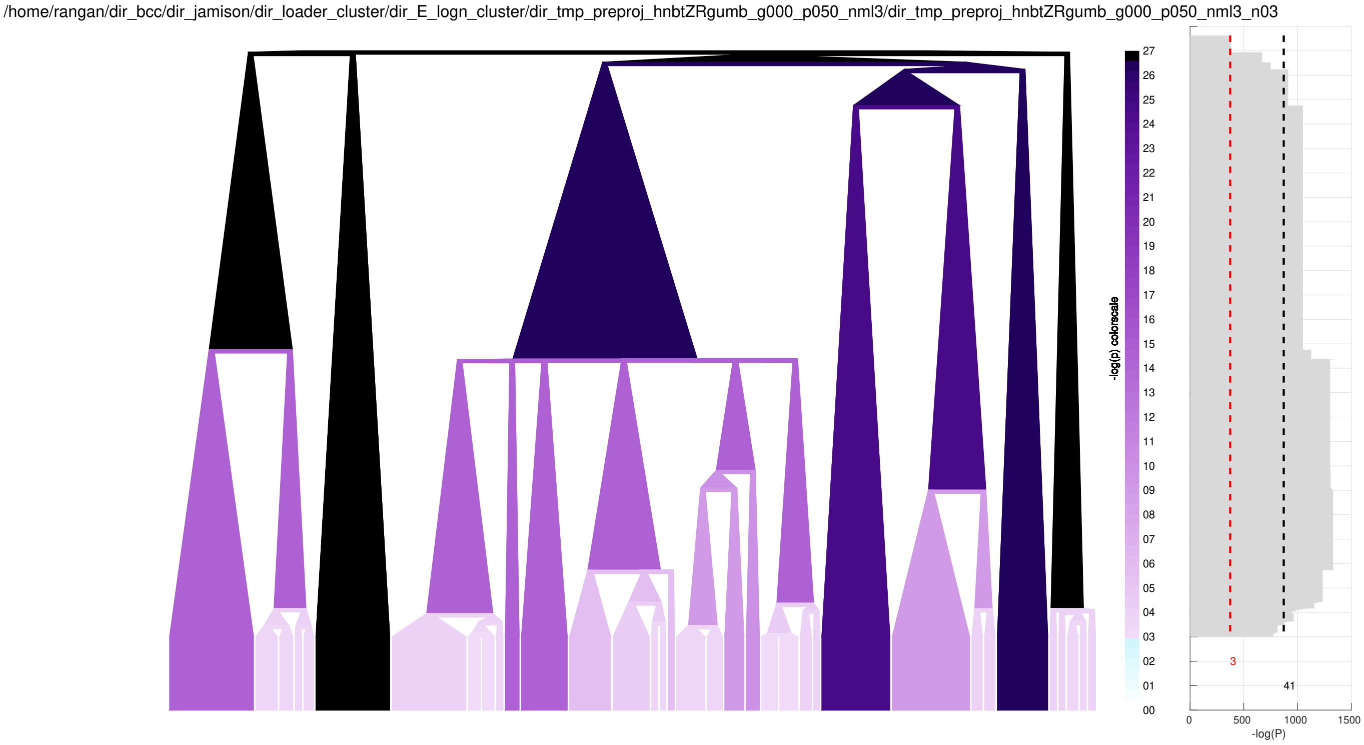


Figure A.6: Here we illustrate the tree and recovered-cluster-quality for the exonic-data described in section A.1.4. The format of this figure is similar to Fig A.2.

refer to the output of the second clustering-algorithm as the disjoint covering

$$\vec{\mathcal{B}} = \{\mathcal{B}[1], \mathcal{B}[2], \ldots, \mathcal{B}[k], \ldots, \mathcal{B}[K]\},$$

where the number of elements in each $\mathcal{B}[k]$ is referred to as $B[k]$. Using this notation, the 'confusion-matrix' $\vec{\mathcal{C}}$ is defined as the array of set-intersections:

$$\mathcal{C}[j,k] = \mathcal{A}[j] \cap \mathcal{B}[k],$$

and we'll refer to the number of elements in each such intersection as the matrix $C[j,k]$. Note that $C$ is constrained: the row- and column-sums of $C$ must equal $\vec{A}$ and $\vec{B}$, respectively. We'll refer to this admissible (convex) set as $\Omega$.

We consider the null-hypothesis that the sample-labels (i.e., cluster-labels for each sample) in $\mathcal{A}$ and/or $\mathcal{B}$ are randomly permuted. With this null-hypothesis, we calculate the 'local' probability $P_0(C)$, which is the probability of drawing a sample from the null-hypothesis with a confusion-matrix $C$:

$$P_0(C) = \frac{\Pi_j A[j]! \cdot \Pi_k B[k]!}{M! \cdot \Pi_{j,k} C[j,k]!}$$

Using this definition for $P_0(C)$, we define a P-value $P$ for the clusterings $\mathcal{A}$ and $\mathcal{B}$:

$$P = \sum_{D \in \Omega \ | \ P_0(D) \leq P_0(C)} P_0(D),$$

where the sum is over all possible confusion-matrices $D \in \Omega$ for which $P_0(D) \leq P_0(C)$. In other words, this P-value is the probability that a sample drawn from the null-hypothesis produces a confusion-matrix that is at least as rare

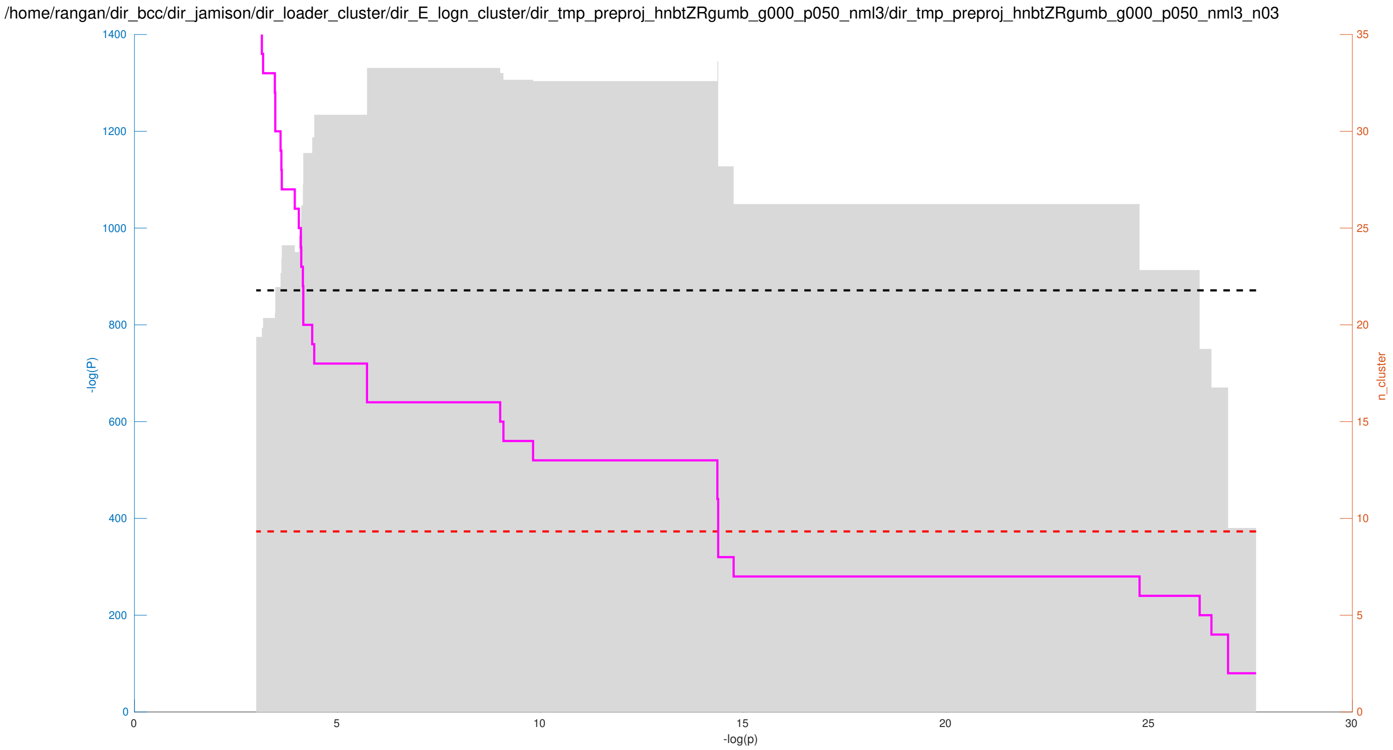


Figure A.7: Here we reorganize the data shown in Fig A.6. We plot the quality $\log(1/P)$ (vertical) as a function of the p-value-threshold (horizontal), reproducing the gray staircase shown on the right of Fig A.6. Additionally, we show the corresponding number of recovered-clusters in magenta. Wide 'plateaus' in the number of recovered-clusters correspond to branches of the tree which persist across a range of p-value-thresholds. These stable plateaus suggest natural cut-points for the p-value-threshold.

/home/rangan/dir_bcc/dir_jamison/dir_loader_cluster/dir_l_logn_cluster/dir_tmp_preproj_hnbtZRgumb_g000_p050_nml3/dir_tmp_preproj_hnbtZRgumb_g000_p050_nml3_n03

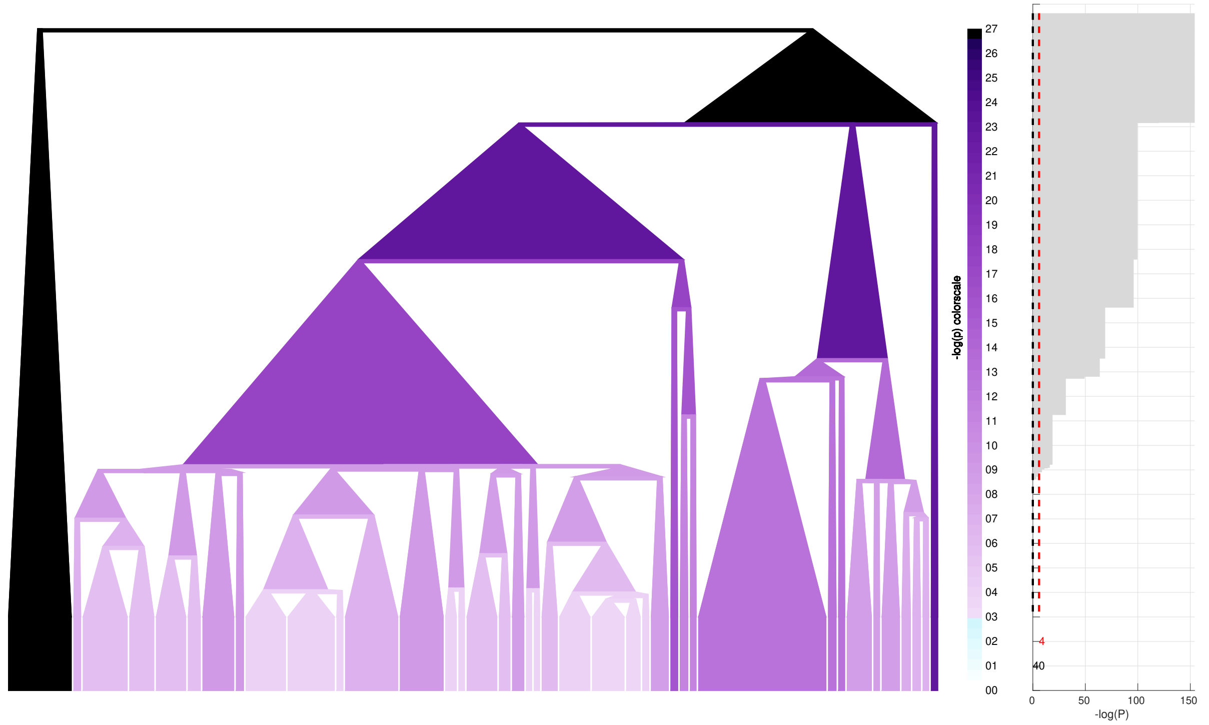


Figure A.8: Here we illustrate the tree and recovered-cluster-quality for the intronic-data described in section A.1.4.

/home/rangan/dir_bcc/dir_jamison/dir_loader_cluster/dir_l_logn_cluster/dir_tmp_preproj_hnbtZRgumb_g000_p050_nml3/dir_tmp_preproj_hnbtZRgumb_g000_p050_nml3_n03

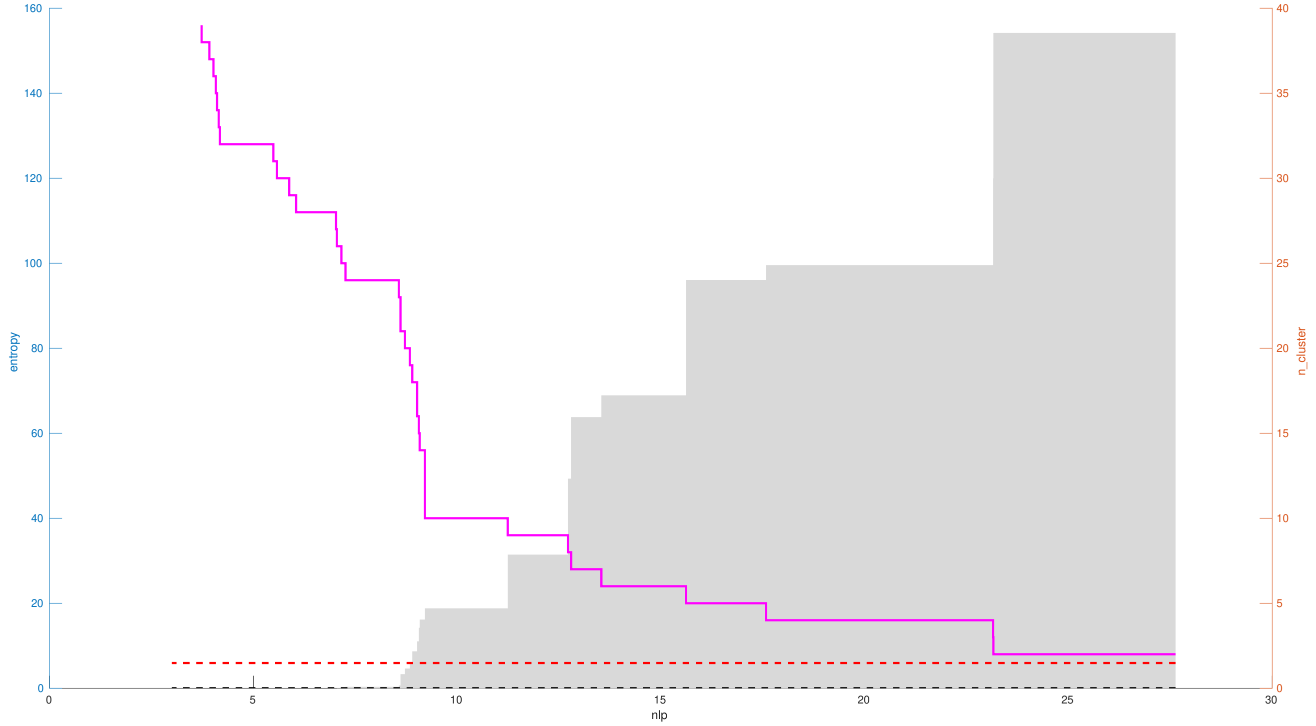


Figure A.9: Here we reorganize the data shown in Fig A.8.

as the confusion-matrix observed between the given clusterings $\mathcal{A}$ and $\mathcal{B}$.

This strategy is related to other strategies for comparing clustering results, such as adjusted-mutual-information [11]. However, we believe that this approach is more easily interpreted, as these P-values are simple probabilities based on a straightforward null-hypothesis.

When $\mathcal{A}$ and $\mathcal{B}$ are sufficiently simple, then $P$ can be calculated directly. For example, if $J$ and $K$ are both equal to 2, then $P$ is simply:

$$P = \sum_{D[1,1]=0,M} P_0\left(\begin{bmatrix} D[1,1] & A[1]-D[1,1] \\ B[1]-D[1,1] & M-A[1]-B[1]+D[1,1] \end{bmatrix}\right),$$

where we define $P_0(D)$ to equal 0 whenever any matrix-entry of $D$ is negative. However, when $\mathcal{A}$ and $\mathcal{B}$ are more complicated (with $J$,$K$ larger than 2), it becomes more difficult to calculate $P$ directly; the sum above now involves a number of terms which grows exponentially with $J$ and $K$. In these situations we approximate $P$ by approximating the function $P_0(D)$ with an anisotropic-gaussian and ignoring the positivity constraints on the entries of $D$ (while retaining the linear constraints on the row- and column-sums of $D$). Using this simple gaussian-approximation, the approximation to the P-value is given by the integral:

$$P \approx \frac{1}{\left(\sqrt{2\pi}\right)^d} \cdot \frac{d\pi^{d/2}}{\Gamma(1+d/2)} \cdot \int_R^\infty \exp(-\frac{1}{2}r^2) \cdot r^{d-1} dr,$$

where $d := JK - J - K + 1$ is the dimensionality of $\Omega$ and $R = \log(1/P_0(C)) + \log(P_0(\vec{A}\vec{B}^\intercal/M))$ is the difference between the negative-log-likelihood of the data $C$ and that of the most likely configuration. Note that this can be solved for analytically using integration-by-parts (see Eq. 313.14 in [12]) producing:

$$P \approx \frac{\Gamma(d/2, R^2/2)}{\Gamma(d/2)}, \tag{A.1}$$

where the numerator references the incomplete gamma-function.

This gaussian-approximation typically provides an overestimate for the true P-value, since it assumes a domain of integration which includes regions where entries of $D$ take on negative values. This conservative estimate is usually acceptable, as we are using it to estimate significance.
