## Appendix B for "Detecting Boolean Asymmetric Relationships with a Loop Counting Technique and its Implications for Analyzing Heterogeneity within Gene Expression Datasets"

### GSEA Results as the Difference Between AD- and WT-Biclusters

All gene pathways are listed in the form: gene pathway name + its p-value.

#### B.1 Gene Set: go_bp_iea

##### B.1.1 Gene Pathways Significant for WT- but not for AD-Biclusters

###### Choroid Plexus

RNA catabolic process 1.100000e-18
SRP dependent cotranslational protein targeting to membrane 6.900000e-22
aromatic compound catabolic process 4.500000e-13
cellular component disassembly 4.200000e-16
cellular macromolecule catabolic process 2.200000e-13
cellular macromolecule localization 5.600000e-12
cellular nitrogen compound catabolic process 4.300000e-13
cellular protein complex disassembly 3.900000e-20
cellular protein localization 5.300000e-12
cotranslational protein targeting to membrane 8.400000e-22
cytoplasmic transport 4.300000e-13
establishment of protein localization to endoplasmic reticulum 9.300000e-22
establishment of protein localization to membrane 8.000000e-19
establishment of protein localization to organelle 1.700000e-15
heterocycle catabolic process 4.300000e-13
interspecies interaction between organisms 2.500000e-13
intracellular protein transport 2.000000e-13
mRNA catabolic process 2.700000e-19
mRNA metabolic process 2.800000e-14
macromolecular complex disassembly 1.800000e-19
macromolecule catabolic process 2.400000e-12

membrane organization 1.800000e-14
multi organism cellular process 1.100000e-13
nuclear transcribed mRNA catabolic process 1.600000e-19
nuclear transcribed mRNA catabolic process nonsense mediated decay 2.000000e-21
nucleobase containing compound catabolic process 2.400000e-13
organic cyclic compound catabolic process 6.600000e-13
protein complex disassembly 1.500000e-19
protein localization to endoplasmic reticulum 3.200000e-21
protein localization to membrane 5.200000e-18
protein localization to organelle 1.500000e-14
protein targeting 3.800000e-15
protein targeting to ER 8.400000e-22
protein targeting to membrane 2.700000e-20
rRNA processing 2.500000e-03
ribonucleoprotein complex biogenesis 2.100000e-05
ribosomal small subunit biogenesis 5.900000e-05
ribosome biogenesis 2.500000e-06
single organism membrane organization 1.200000e-14
symbiosis encompassing mutualism through parasitism 2.500000e-13
translation 2.500000e-15
translational elongation 5.600000e-22
translational initiation 2.600000e-20
translational termination 1.100000e-22
viral gene expression 3.900000e-20
viral life cycle 5.600000e-18
viral process 1.000000e-13
viral transcription 1.900000e-20

**GABAergic Neurons**

ATP synthesis coupled proton transport 2.600000e-03
RNA catabolic process 5.100000e-22
SRP dependent cotranslational protein targeting to membrane 1.800000e-30
aromatic compound catabolic process 2.900000e-12
cellular component disassembly 3.000000e-18
cellular macromolecule catabolic process 6.900000e-14
cellular macromolecule localization 5.700000e-13
cellular nitrogen compound catabolic process 2.600000e-12
cellular protein complex disassembly 6.300000e-25
cellular protein localization 5.100000e-13
cotranslational protein targeting to membrane 2.800000e-30
cytoplasmic transport 2.400000e-14
energy coupled proton transport down electrochemical gradient 2.600000e-03
establishment of protein localization to endoplasmic reticulum 3.500000e-30

establishment of protein localization to membrane 7.200000e-24
establishment of protein localization to organelle 1.400000e-19
heterocycle catabolic process 2.700000e-12
interaction with host 1.900000e-03
interspecies interaction between organisms 2.000000e-19
intracellular protein transport 5.800000e-14
mRNA catabolic process 3.200000e-23
mRNA metabolic process 6.500000e-17
macromolecular complex disassembly 1.400000e-23
macromolecule catabolic process 8.200000e-12
membrane organization 4.100000e-16
modification by symbiont of host morphology or physiology 9.900000e-04
modification of morphology or physiology of other organism 1.700000e-03
modification of morphology or physiology of other organism involved in symbiotic interaction 1.400000e-03
modulation by virus of host morphology or physiology 8.800000e-04
multi organism cellular process 2.200000e-20
negative regulation of RNA splicing 2.600000e-03
nuclear transcribed mRNA catabolic process 1.100000e-23
nuclear transcribed mRNA catabolic process nonsense mediated decay 1.500000e-27
nucleobase containing compound catabolic process 8.900000e-13
organic cyclic compound catabolic process 6.100000e-12
protein complex disassembly 9.900000e-24
protein localization to endoplasmic reticulum 5.100000e-29
protein localization to membrane 3.600000e-22
protein localization to organelle 1.800000e-17
protein targeting 1.500000e-17
protein targeting to ER 2.800000e-30
protein targeting to membrane 5.400000e-27
ribosomal small subunit biogenesis 3.800000e-05
ribosome biogenesis 3.500000e-04
single organism membrane organization 1.800000e-16
symbiosis encompassing mutualism through parasitism 2.000000e-19
translation 3.400000e-19
translational elongation 1.100000e-30
translational initiation 2.700000e-25
translational termination 3.400000e-30
viral gene expression 1.200000e-26
viral life cycle 3.400000e-25
viral process 2.000000e-20
viral transcription 1.500000e-25
virion assembly 1.800000e-03

**Radial Glial Cells**

Fc gamma receptor signaling pathway 9.100000e-05
Fc gamma receptor signaling pathway involved in phagocytosis 9.100000e-05
Fc receptor mediated stimulatory signaling pathway 9.900000e-05
Fc receptor signaling pathway 8.300000e-04
G2 M transition of mitotic cell cycle 3.400000e-04
activation of immune response 2.300000e-03
axon guidance 2.100000e-03
cell type specific apoptotic process 1.400000e-03
cellular protein complex assembly 1.700000e-03
cellular response to interleukin 4 1.000000e-05
immune response activating cell surface receptor signaling pathway 6.200000e-04
immune response activating signal transduction 1.700000e-03
immune response regulating cell surface receptor signaling pathway 1.600000e-03
immune response regulating cell surface receptor signaling pathway involved in phagocytosis 9.100000e-05
neuron projection guidance 2.100000e-03
nuclear import 8.600000e-04
nuclear transport 2.200000e-03
nucleocytoplasmic transport 2.200000e-03
phagocytosis 4.200000e-04
positive regulation of intracellular protein transport 1.900000e-04
positive regulation of intracellular transport 3.600000e-04
positive regulation of protein import into nucleus translocation 2.100000e-06
positive regulation of protein transport 5.600000e-04
protein folding 3.400000e-06
protein import 1.100000e-03
protein import into nucleus 8.400000e-04
protein import into nucleus translocation 2.900000e-05
protein localization to nucleus 1.200000e-03
protein targeting to nucleus 8.400000e-04
regulation of anatomical structure size 1.500000e-03
regulation of cell size 1.000000e-04
regulation of cellular component size 5.600000e-04
regulation of establishment of protein localization 2.100000e-03
regulation of intracellular protein transport 5.600000e-04
regulation of intracellular transport 1.700000e-03
regulation of nucleocytoplasmic transport 5.000000e-04
regulation of protein import into nucleus 3.200000e-04
regulation of protein import into nucleus translocation 5.100000e-06
regulation of protein localization to nucleus 3.400000e-04
regulation of protein transport 1.600000e-03
response to interleukin 4 1.300000e-05
response to osmotic stress 5.000000e-05

response to salt stress 8.700000e-06
response to topologically incorrect protein 3.500000e-04
response to unfolded protein 3.200000e-04

##### B.1.2 Gene Pathways Significant for AD- but not for WT-Biclusters

**Astroglial Cells**

ATP biosynthetic process 8.700000e-05
ATP hydrolysis coupled proton transport 6.900000e-03
ATP metabolic process 4.000000e-03
ATP synthesis coupled electron transport 3.800000e-04
ATP synthesis coupled proton transport 2.300000e-06
DNA damage response signal transduction by p53 class mediator 2.900000e-03
DNA damage response signal transduction by p53 class mediator resulting in cell cycle arrest 1.900000e-03
Fc gamma receptor signaling pathway 3.600000e-04
Fc gamma receptor signaling pathway involved in phagocytosis 3.600000e-04
Fc receptor mediated stimulatory signaling pathway 4.600000e-04
Fc receptor signaling pathway 1.400000e-04
G1 DNA damage checkpoint 2.900000e-03
MyD88 dependent toll like receptor signaling pathway 5.100000e-03
Notch receptor processing 1.300000e-04
RNA catabolic process 1.300000e-04
RNA catabolic process 6.000000e-16
RNA catabolic process 9.500000e-61
RNA processing 5.800000e-07
RNA splicing 5.000000e-04
RNA splicing 7.300000e-06
RNA splicing via transesterification reactions 1.400000e-06
RNA splicing via transesterification reactions with bulged adenosine as nucleophile 1.000000e-06
RNA stabilization 4.500000e-03
SRP dependent cotranslational protein targeting to membrane 2.700000e-87
SRP dependent cotranslational protein targeting to membrane 4.300000e-06
SRP dependent cotranslational protein targeting to membrane 8.900000e-20
de novo posttranslational protein folding 4.400000e-05
de novo protein folding 6.900000e-05
acid amino acid ligase activity 1.800000e-03
activation of immune response 9.200000e-03
adipose tissue development 2.700000e-03
anaphase promoting complex dependent proteasomal ubiquitin dependent protein catabolic process 7.000000e-04
antigen processing and presentation of exogenous antigen 6.800000e-03
antigen processing and presentation of exogenous peptide antigen 6.300000e-03
antigen processing and presentation of exogenous peptide antigen via MHC class I 7.400000e-05
antigen processing and presentation of exogenous peptide antigen via MHC class I TAP dependent 5.200000e-05

antigen processing and presentation of peptide antigen 1.000000e-02
antigen processing and presentation of peptide antigen via MHC class I 3.400000e-04
apoptotic mitochondrial changes 1.300000e-03
apoptotic signaling pathway 5.900000e-03
aromatic compound catabolic process 1.200000e-10
aromatic compound catabolic process 1.800000e-03
aromatic compound catabolic process 2.300000e-35
axonogenesis 4.900000e-03
calcium ion dependent exocytosis 5.000000e-03
cell division 7.900000e-04
cell part morphogenesis 3.500000e-03
cell projection morphogenesis 3.100000e-03
cell projection organization 3.900000e-03
cellular component disassembly 1.900000e-46
cellular component disassembly 2.400000e-04
cellular component disassembly 4.100000e-16
cellular macromolecular complex assembly 3.200000e-06
cellular macromolecule catabolic process 2.400000e-39
cellular macromolecule catabolic process 4.900000e-11
cellular macromolecule localization 1.300000e-32
cellular macromolecule localization 5.800000e-05
cellular macromolecule localization 8.800000e-12
cellular nitrogen compound catabolic process 1.100000e-10
cellular nitrogen compound catabolic process 1.600000e-35
cellular nitrogen compound catabolic process 1.700000e-03
cellular protein complex assembly 8.700000e-03
cellular protein complex disassembly 1.400000e-06
cellular protein complex disassembly 1.400000e-66
cellular protein complex disassembly 9.800000e-22
cellular protein localization 5.500000e-05
cellular protein localization 8.200000e-12
cellular protein localization 9.600000e-33
cellular respiration 4.800000e-03
cellular respiration 6.200000e-09
cellular response to decreased oxygen levels 6.200000e-03
cellular response to hypoxia 6.200000e-03
cellular response to interleukin 4 4.000000e-03
cellular response to oxygen levels 8.600000e-03
cellular response to superoxide 7.700000e-03
cotranslational protein targeting to membrane 1.100000e-19
cotranslational protein targeting to membrane 1.500000e-86
cotranslational protein targeting to membrane 4.700000e-06
cytoplasmic pattern recognition receptor signaling pathway 2.400000e-03

cytoplasmic transport 1.100000e-10
cytoplasmic transport 3.100000e-04
cytoplasmic transport 8.200000e-42
electron transport chain 1.600000e-03
electron transport chain 3.800000e-11
endoplasmic reticulum organization 1.600000e-03
energy coupled proton transmembrane transport against electrochemical gradient 6.900000e-03
energy coupled proton transport down electrochemical gradient 2.300000e-06
energy derivation by oxidation of organic compounds 2.900000e-06
establishment of protein localization to endoplasmic reticulum 1.300000e-19
establishment of protein localization to endoplasmic reticulum 3.400000e-86
establishment of protein localization to endoplasmic reticulum 5.000000e-06
establishment of protein localization to membrane 3.100000e-68
establishment of protein localization to membrane 4.200000e-16
establishment of protein localization to membrane 7.800000e-06
establishment of protein localization to organelle 2.800000e-15
establishment of protein localization to organelle 6.200000e-06
establishment of protein localization to organelle 8.300000e-52
female meiosis 8.000000e-04
filopodium assembly 3.700000e-03
forebrain generation of neurons 4.000000e-03
forebrain neuron development 6.300000e-04
forebrain neuron differentiation 1.900000e-03
generation of neurons 4.400000e-04
generation of precursor metabolites and energy 8.600000e-09
glutamate secretion 2.300000e-03
glycolysis 7.800000e-03
heterocycle catabolic process 1.100000e-10
heterocycle catabolic process 1.700000e-03
heterocycle catabolic process 1.800000e-35
hydrogen peroxide metabolic process 9.900000e-03
hydrogen transport 2.400000e-04
hypothalamus development 1.400000e-03
immune response activating cell surface receptor signaling pathway 1.900000e-04
immune response activating signal transduction 3.000000e-03
immune response regulating cell surface receptor signaling pathway 2.300000e-03
immune response regulating cell surface receptor signaling pathway involved in phagocytosis 3.600000e-04
immune response regulating signaling pathway 9.200000e-03
interaction with host 2.700000e-03
interspecies interaction between organisms 1.300000e-03
interspecies interaction between organisms 2.600000e-12
interspecies interaction between organisms 8.200000e-47
intracellular protein transport 1.400000e-39

intracellular protein transport 1.900000e-12
intracellular protein transport 3.200000e-06
intracellular signal transduction involved in G1 DNA damage checkpoint 1.900000e-03
intrinsic apoptotic signaling pathway 6.800000e-04
ion transmembrane transport 5.800000e-03
ligase activity 2.000000e-03
ligase activity forming carbon nitrogen bonds 1.800000e-03
limbic system development 1.000000e-02
localization within membrane 2.900000e-03
localization within membrane 4.700000e-03
mRNA catabolic process 1.200000e-16
mRNA catabolic process 1.400000e-64
mRNA catabolic process 7.000000e-05
mRNA metabolic process 3.000000e-56
mRNA metabolic process 3.600000e-12
mRNA metabolic process 4.600000e-05
mRNA processing 1.200000e-04
mRNA splicing via spliceosome 1.000000e-06
mRNA stabilization 4.500000e-03
macromolecular complex disassembly 3.400000e-06
macromolecular complex disassembly 4.900000e-64
macromolecular complex disassembly 8.500000e-21
macromolecule catabolic process 4.500000e-03
macromolecule catabolic process 6.800000e-34
macromolecule catabolic process 9.500000e-10
male meiosis 8.300000e-03
male meiosis I 1.400000e-03
membrane organization 1.000000e-42
membrane organization 3.400000e-05
membrane organization 7.400000e-14
mitochondrial ATP synthesis coupled electron transport 3.800000e-04
mitochondrial ATP synthesis coupled proton transport 8.000000e-04
mitochondrial outer membrane permeabilization 2.200000e-03
mitochondrial outer membrane permeabilization 3.700000e-03
mitochondrial transport 3.500000e-03
mitochondrial transport 5.200000e-03
mitotic DNA damage checkpoint 3.700000e-03
mitotic DNA integrity checkpoint 4.300000e-03
mitotic G1 DNA damage checkpoint 2.700000e-03
modification by symbiont of host morphology or physiology 3.100000e-03
modification of morphology or physiology of other organism 6.500000e-03
modification of morphology or physiology of other organism involved in symbiotic interaction 4.900000e-03
modulation by virus of host morphology or physiology 2.600000e-03

multi organism cellular process 2.800000e-48
multi organism cellular process 7.600000e-04
multi organism cellular process 8.000000e-13
ncRNA metabolic process 4.200000e-03
ncRNA processing 4.900000e-03
negative regulation of ERBB signaling pathway 2.000000e-03
negative regulation of G1 S transition of mitotic cell cycle 6.500000e-03
negative regulation of RNA metabolic process 8.300000e-05
negative regulation of RNA splicing 1.400000e-03
negative regulation of apoptotic process 3.400000e-04
negative regulation of biosynthetic process 3.200000e-04
negative regulation of cell death 4.700000e-04
negative regulation of cellular biosynthetic process 2.900000e-04
negative regulation of cellular component organization 3.300000e-03
negative regulation of cellular macromolecule biosynthetic process 1.400000e-04
negative regulation of cellular protein metabolic process 1.600000e-03
negative regulation of cellular response to growth factor stimulus 4.800000e-03
negative regulation of cytokine production 9.200000e-03
negative regulation of endocytosis 5.100000e-03
negative regulation of epidermal growth factor receptor signaling pathway 2.000000e-03
negative regulation of gene expression 2.200000e-05
negative regulation of ligase activity 2.600000e-03
negative regulation of macromolecule biosynthetic process 2.000000e-04
negative regulation of nitrogen compound metabolic process 2.000000e-04
negative regulation of nucleobase containing compound metabolic process 1.800000e-04
negative regulation of programmed cell death 3.600000e-04
negative regulation of protein metabolic process 5.600000e-03
negative regulation of protein polymerization 3.000000e-03
negative regulation of protein ubiquitination 1.000000e-02
negative regulation of transcription DNA dependent 5.900000e-05
negative regulation of transcription from RNA polymerase II promoter 8.800000e-04
negative regulation of transforming growth factor beta receptor signaling pathway 1.200000e-03
negative regulation of translation 6.500000e-03
negative regulation of transmembrane receptor protein serine threonine kinase signaling pathway 5.900000e-03
negative regulation of type I interferon production 1.200000e-03
negative regulation of ubiquitin protein ligase activity 2.600000e-03
negative regulation of ubiquitin protein ligase activity involved in mitotic cell cycle 2.000000e-03
neuron development 1.800000e-03
neuron differentiation 1.300000e-03
neuron projection development 7.400000e-04
neuron projection morphogenesis 1.300000e-03
nuclear transcribed mRNA catabolic process 4.300000e-66
nuclear transcribed mRNA catabolic process 5.500000e-05

nuclear transcribed mRNA catabolic process 6.100000e-17

nuclear transcribed mRNA catabolic process nonsense mediated decay 2.600000e-77

nuclear transcribed mRNA catabolic process nonsense mediated decay 3.300000e-19

nuclear transcribed mRNA catabolic process nonsense mediated decay 7.100000e-06

nuclear transport 7.900000e-03

nuclease activity 5.700000e-04

nucleic acid phosphodiester bond hydrolysis 2.300000e-04

nucleobase containing compound catabolic process 1.200000e-03

nucleobase containing compound catabolic process 5.500000e-11

nucleobase containing compound catabolic process 5.500000e-37

nucleocytoplasmic transport 7.100000e-03

nucleoside monophosphate biosynthetic process 3.300000e-03

nucleoside triphosphate biosynthetic process 1.500000e-03

nucleotide binding domain leucine rich repeat containing receptor signaling pathway 5.100000e-04

nucleotide binding oligomerization domain containing signaling pathway 8.700000e-04

organic cyclic compound catabolic process 1.900000e-10

organic cyclic compound catabolic process 2.200000e-03

organic cyclic compound catabolic process 2.300000e-34

oxidation reduction process 4.400000e-04

oxidative phosphorylation 1.100000e-03

phagocytosis 1.000000e-03

positive regulation of apoptotic process 1.200000e-02

positive regulation of cell cycle arrest 4.300000e-03

positive regulation of cell projection organization 3.200000e-03

positive regulation of cellular component organization 9.700000e-04

positive regulation of ligase activity 1.000000e-03

positive regulation of mitochondrial membrane permeability 2.500000e-03

positive regulation of mitochondrial membrane permeability 4.100000e-03

positive regulation of mitochondrial membrane permeability involved in apoptotic process 2.300000e-03

positive regulation of mitochondrial membrane permeability involved in apoptotic process 3.900000e-03

positive regulation of mitochondrion organization 1.700000e-04

positive regulation of mitochondrion organization 3.100000e-03

positive regulation of nuclease activity 4.400000e-04

positive regulation of organelle organization 1.900000e-03

positive regulation of protein import into nucleus translocation 9.300000e-03

positive regulation of protein insertion into mitochondrial membrane involved in apoptotic signaling pathway 1.600000e-03

positive regulation of protein insertion into mitochondrial membrane involved in apoptotic signaling pathway 9.300000e-04

positive regulation of protein ubiquitination 9.800000e-03

positive regulation of translation 3.200000e-03

positive regulation of type I interferon production 2.600000e-03

positive regulation of ubiquitin protein ligase activity 8.500000e-04

positive regulation of ubiquitin protein ligase activity involved in mitotic cell cycle 3.900000e-04
posttranscriptional regulation of gene expression 2.400000e-07
proteasomal protein catabolic process 1.800000e-03
proteasomal ubiquitin dependent protein catabolic process 1.200000e-03
protein complex disassembly 1.600000e-64
protein complex disassembly 3.100000e-06
protein complex disassembly 6.600000e-21
protein folding 1.100000e-05
protein folding 7.700000e-05
protein insertion into membrane 2.000000e-03
protein insertion into membrane 3.300000e-03
protein insertion into mitochondrial membrane 1.100000e-03
protein insertion into mitochondrial membrane 1.800000e-03
protein insertion into mitochondrial membrane involved in apoptotic signaling pathway 1.000000e-03
protein insertion into mitochondrial membrane involved in apoptotic signaling pathway 1.700000e-03
protein localization to endoplasmic reticulum 5.600000e-19
protein localization to endoplasmic reticulum 6.000000e-82
protein localization to endoplasmic reticulum 8.800000e-06
protein localization to membrane 2.200000e-05
protein localization to membrane 3.800000e-15
protein localization to membrane 9.700000e-63
protein localization to organelle 3.000000e-05
protein localization to organelle 5.700000e-14
protein localization to organelle 8.100000e-47
protein polymerization 3.900000e-03
protein refolding 4.200000e-04
protein targeting 1.100000e-05
protein targeting 1.600000e-49
protein targeting 3.000000e-13
protein targeting to ER 1.100000e-19
protein targeting to ER 1.500000e-86
protein targeting to ER 4.700000e-06
protein targeting to membrane 2.400000e-05
protein targeting to membrane 5.500000e-75
protein targeting to membrane 7.400000e-18
proton transport 2.000000e-04
purine nucleoside biosynthetic process 4.600000e-03
purine nucleoside monophosphate biosynthetic process 1.000000e-03
purine nucleoside monophosphate metabolic process 1.200000e-02
purine nucleoside triphosphate biosynthetic process 3.800000e-04
purine ribonucleoside biosynthetic process 4.600000e-03
purine ribonucleoside monophosphate biosynthetic process 1.000000e-03
purine ribonucleoside monophosphate metabolic process 1.100000e-02

purine ribonucleoside triphosphate biosynthetic process 3.400000e-04
rRNA metabolic process 7.700000e-05
rRNA metabolic process 9.500000e-04
rRNA processing 5.000000e-05
rRNA processing 7.900000e-04
regulation of DNA dependent transcription in response to stress 1.900000e-04
regulation of ERBB signaling pathway 9.300000e-03
regulation of RNA splicing 9.900000e-03
regulation of RNA stability 1.500000e-03
regulation of cell cycle arrest 8.200000e-03
regulation of cell development 4.400000e-03
regulation of cell projection assembly 5.200000e-03
regulation of cell projection organization 2.400000e-03
regulation of epidermal growth factor receptor signaling pathway 8.800000e-03
regulation of establishment of protein localization 1.300000e-03
regulation of filopodium assembly 1.400000e-03
regulation of intracellular transport 2.400000e-03
regulation of intracellular transport 7.900000e-04
regulation of lamellipodium assembly 9.300000e-03
regulation of ligase activity 2.000000e-03
regulation of mRNA stability 1.200000e-03
regulation of mitochondrial membrane permeability 3.400000e-03
regulation of mitochondrial membrane permeability involved in apoptotic process 2.600000e-03
regulation of mitochondrial membrane permeability involved in apoptotic process 4.300000e-03
regulation of mitochondrial outer membrane permeabilization 1.800000e-03
regulation of mitochondrial outer membrane permeabilization 2.900000e-03
regulation of mitochondrion organization 6.600000e-04
regulation of nuclease activity 5.700000e-04
regulation of organelle organization 1.700000e-03
regulation of organelle organization 2.900000e-03
regulation of protein insertion into mitochondrial membrane involved in apoptotic signaling pathway 1.600000e-03
regulation of protein insertion into mitochondrial membrane involved in apoptotic signaling pathway 9.300000e-04
regulation of protein localization 2.600000e-03
regulation of transcription from RNA polymerase II promoter in response to hypoxia 2.700000e-05
regulation of transcription from RNA polymerase II promoter in response to stress 1.200000e-04
regulation of transforming growth factor beta receptor signaling pathway 6.200000e-03
regulation of translation 9.900000e-08
regulation of translational elongation 7.700000e-03
regulation of translational initiation 1.100000e-05
regulation of ubiquitin protein ligase activity 1.700000e-03
regulation of ubiquitin protein ligase activity involved in mitotic cell cycle 6.100000e-04

removal of superoxide radicals 7.700000e-03
respiratory electron transport chain 1.500000e-03
respiratory electron transport chain 2.800000e-11
response to inorganic substance 7.500000e-04
response to interleukin 4 5.600000e-03
response to oxidative stress 2.200000e-04
response to reactive oxygen species 1.800000e-04
ribonucleoprotein complex assembly 2.800000e-05
ribonucleoprotein complex biogenesis 1.500000e-08
ribonucleoprotein complex biogenesis 8.700000e-04
ribonucleoprotein complex subunit organization 3.200000e-05
ribonucleoside biosynthetic process 9.400000e-03
ribonucleoside monophosphate biosynthetic process 2.700000e-03
ribonucleoside triphosphate biosynthetic process 6.100000e-04
ribosomal large subunit biogenesis 1.900000e-07
ribosomal small subunit biogenesis 2.600000e-08
ribosomal small subunit biogenesis 3.600000e-04
ribosome assembly 6.800000e-05
ribosome biogenesis 2.100000e-03
ribosome biogenesis 2.900000e-07
signal transduction by p53 class mediator 2.000000e-03
signal transduction in response to DNA damage 4.500000e-03
signal transduction involved in DNA damage checkpoint 2.000000e-03
signal transduction involved in DNA integrity checkpoint 2.000000e-03
signal transduction involved in cell cycle checkpoint 2.100000e-03
signal transduction involved in mitotic DNA damage checkpoint 1.900000e-03
signal transduction involved in mitotic DNA integrity checkpoint 1.900000e-03
signal transduction involved in mitotic G1 DNA damage checkpoint 1.900000e-03
signal transduction involved in mitotic cell cycle checkpoint 1.900000e-03
single organism membrane organization 2.600000e-05
single organism membrane organization 4.300000e-14
single organism membrane organization 8.400000e-44
small conjugating protein ligase activity 1.700000e-03
symbiosis encompassing mutualism through parasitism 1.300000e-03
symbiosis encompassing mutualism through parasitism 2.600000e-12
symbiosis encompassing mutualism through parasitism 8.200000e-47
synaptic vesicle exocytosis 2.400000e-03
toll like receptor 10 signaling pathway 1.200000e-02
translation 4.900000e-15
translation 7.800000e-05
translation 8.600000e-63
translational elongation 3.900000e-06
translational elongation 4.400000e-22

translational elongation 4.700000e-88
translational initiation 1.100000e-06
translational initiation 6.000000e-81
translational initiation 6.800000e-18
translational termination 1.800000e-06
translational termination 7.500000e-85
translational termination 9.900000e-21
ubiquitin protein ligase activity 1.700000e-03
viral gene expression 1.100000e-17
viral gene expression 2.900000e-05
viral gene expression 4.800000e-72
viral life cycle 2.800000e-04
viral life cycle 4.200000e-15
viral life cycle 6.300000e-61
viral process 2.200000e-48
viral process 7.400000e-04
viral process 7.700000e-13
viral protein processing 1.500000e-05
viral transcription 1.800000e-70
viral transcription 2.000000e-05
viral transcription 4.800000e-18
virion assembly 7.800000e-07

**Oligodendrocyte Progenitor Cells**

RNA catabolic process 8.000000e-07
SRP dependent cotranslational protein targeting to membrane 2.300000e-08
aromatic compound catabolic process 3.800000e-04
cellular component disassembly 1.400000e-05
cellular macromolecule catabolic process 1.800000e-05
cellular macromolecule localization 1.200000e-03
cellular nitrogen compound catabolic process 3.700000e-04
cellular protein complex disassembly 1.600000e-07
cellular protein localization 1.200000e-03
cotranslational protein targeting to membrane 2.500000e-08
cytoplasmic transport 3.700000e-04
establishment of protein localization to endoplasmic reticulum 2.600000e-08
establishment of protein localization to membrane 6.900000e-07
establishment of protein localization to organelle 2.800000e-05
heterocycle catabolic process 3.700000e-04
interspecies interaction between organisms 1.900000e-05
intracellular protein transport 2.600000e-04
mRNA catabolic process 4.100000e-07
mRNA metabolic process 2.100000e-07

macromolecular complex disassembly 3.400000e-07
macromolecule catabolic process 7.000000e-05
membrane organization 8.600000e-05
multi organism cellular process 1.200000e-05
nuclear transcribed mRNA catabolic process 3.200000e-07
nuclear transcribed mRNA catabolic process nonsense mediated decay 3.800000e-08
nucleobase containing compound catabolic process 2.900000e-04
organic cyclic compound catabolic process 4.600000e-04
protein complex disassembly 3.100000e-07
protein localization to endoplasmic reticulum 4.700000e-08
protein localization to membrane 1.700000e-06
protein localization to organelle 7.800000e-05
protein targeting 4.000000e-05
protein targeting to ER 2.500000e-08
protein targeting to membrane 1.300000e-07
single organism membrane organization 7.100000e-05
symbiosis encompassing mutualism through parasitism 1.900000e-05
translation 1.400000e-06
translational elongation 2.000000e-08
translational initiation 1.300000e-07
translational termination 9.400000e-09
viral gene expression 2.100000e-09
viral life cycle 3.900000e-08
viral process 1.200000e-05
viral transcription 1.100000e-07

**Ventral Progenitors**

RNA catabolic process 2.600000e-22
RNA catabolic process 7.000000e-17
RNA processing 1.700000e-06
RNA splicing 3.800000e-08
RNA splicing via transesterification reactions 1.100000e-07
RNA splicing via transesterification reactions with bulged adenosine as nucleophile 8.600000e-08
SRP dependent cotranslational protein targeting to membrane 3.500000e-26
SRP dependent cotranslational protein targeting to membrane 9.400000e-20
de novo posttranslational protein folding 1.200000e-03
de novo protein folding 1.600000e-03
apoptotic mitochondrial changes 6.200000e-04
apoptotic signaling pathway 1.400000e-03
aromatic compound catabolic process 1.500000e-15
aromatic compound catabolic process 7.800000e-12
cellular component disassembly 1.500000e-14
cellular component disassembly 3.400000e-19

cellular macromolecular complex assembly 2.300000e-03
cellular macromolecule catabolic process 4.100000e-12
cellular macromolecule catabolic process 6.500000e-16
cellular macromolecule localization 3.100000e-14
cellular macromolecule localization 7.500000e-11
cellular nitrogen compound catabolic process 1.400000e-15
cellular nitrogen compound catabolic process 7.400000e-12
cellular protein complex disassembly 3.500000e-18
cellular protein complex disassembly 4.700000e-24
cellular protein localization 2.900000e-14
cellular protein localization 7.200000e-11
cellular respiration 3.800000e-04
cotranslational protein targeting to membrane 1.100000e-19
cotranslational protein targeting to membrane 4.600000e-26
cytoplasmic transport 1.400000e-15
cytoplasmic transport 7.500000e-12
electron transport chain 6.500000e-05
energy derivation by oxidation of organic compounds 2.000000e-03
establishment of protein localization to endoplasmic reticulum 1.200000e-19
establishment of protein localization to endoplasmic reticulum 5.100000e-26
establishment of protein localization to membrane 1.800000e-22
establishment of protein localization to membrane 5.300000e-17
establishment of protein localization to organelle 1.800000e-18
establishment of protein localization to organelle 2.600000e-03
establishment of protein localization to organelle 5.200000e-14
establishment of viral latency 8.500000e-04
generation of precursor metabolites and energy 1.400000e-03
heterocycle catabolic process 1.400000e-15
heterocycle catabolic process 7.500000e-12
hydrogen peroxide metabolic process 1.300000e-05
hydrogen transport 3.000000e-03
interspecies interaction between organisms 4.600000e-12
interspecies interaction between organisms 7.400000e-16
intracellular protein transport 2.900000e-03
intracellular protein transport 3.800000e-12
intracellular protein transport 5.800000e-16
intrinsic apoptotic signaling pathway 9.400000e-04
localization within membrane 3.400000e-05
mRNA catabolic process 2.000000e-17
mRNA catabolic process 4.900000e-23
mRNA metabolic process 1.300000e-08
mRNA metabolic process 5.200000e-17
mRNA metabolic process 6.300000e-13

mRNA processing 4.500000e-09

mRNA splicing via spliceosome 8.600000e-08

macromolecular complex disassembly 1.400000e-17

macromolecular complex disassembly 3.000000e-23

macromolecule catabolic process 1.100000e-14

macromolecule catabolic process 3.500000e-11

membrane organization 3.200000e-17

membrane organization 4.400000e-13

mitochondrial membrane organization 3.100000e-03

mitochondrial outer membrane permeabilization 5.800000e-04

multi organism cellular process 2.100000e-12

multi organism cellular process 2.600000e-16

ncRNA metabolic process 1.900000e-03

ncRNA processing 7.100000e-04

nuclear transcribed mRNA catabolic process 1.300000e-17

nuclear transcribed mRNA catabolic process 2.600000e-23

nuclear transcribed mRNA catabolic process nonsense mediated decay 1.300000e-25

nuclear transcribed mRNA catabolic process nonsense mediated decay 2.500000e-19

nucleobase containing compound catabolic process 4.500000e-12

nucleobase containing compound catabolic process 7.200000e-16

organic cyclic compound catabolic process 1.100000e-11

organic cyclic compound catabolic process 2.400000e-15

positive regulation of cell projection organization 2.000000e-03

positive regulation of cellular component organization 2.400000e-04

positive regulation of mitochondrial membrane permeability 6.800000e-04

positive regulation of mitochondrial membrane permeability involved in apoptotic process 6.300000e-04

positive regulation of mitochondrion organization 4.100000e-05

positive regulation of organelle organization 3.900000e-04

positive regulation of protein insertion into mitochondrial membrane involved in apoptotic signaling pathway 1.600000e-04

posttranscriptional regulation of gene expression 2.000000e-05

protein complex disassembly 1.200000e-17

protein complex disassembly 2.400000e-23

protein folding 1.500000e-03

protein insertion into membrane 4.900000e-04

protein insertion into mitochondrial membrane 2.000000e-04

protein insertion into mitochondrial membrane involved in apoptotic signaling pathway 1.800000e-04

protein localization to endoplasmic reticulum 2.300000e-25

protein localization to endoplasmic reticulum 3.700000e-19

protein localization to membrane 1.700000e-21

protein localization to membrane 2.800000e-16

protein localization to organelle 2.200000e-03

protein localization to organelle 2.600000e-17

protein localization to organelle 3.700000e-13
protein targeting 1.100000e-13
protein targeting 4.800000e-18
protein targeting to ER 1.100000e-19
protein targeting to ER 4.600000e-26
protein targeting to membrane 2.600000e-18
protein targeting to membrane 3.100000e-24
proton transport 2.700000e-03
rRNA metabolic process 1.100000e-04
rRNA processing 9.200000e-05
reactive oxygen species metabolic process 1.500000e-04
regulation of apoptotic signaling pathway 1.300000e-04
regulation of cell projection assembly 9.700000e-04
regulation of cellular localization 2.800000e-03
regulation of establishment of protein localization 7.300000e-04
regulation of intracellular transport 5.300000e-05
regulation of mitochondrial membrane permeability 1.100000e-03
regulation of mitochondrial membrane permeability involved in apoptotic process 7.300000e-04
regulation of mitochondrial outer membrane permeabilization 4.100000e-04
regulation of mitochondrion organization 2.600000e-04
regulation of organelle organization 5.200000e-04
regulation of protein insertion into mitochondrial membrane involved in apoptotic signaling pathway 1.600000e-04
regulation of protein localization 1.800000e-03
regulation of translation 2.000000e-03
regulation of translational initiation 1.200000e-04
regulation of transport 2.600000e-04
respiratory electron transport chain 5.800000e-05
response to inorganic substance 2.400000e-03
response to superoxide 1.700000e-03
ribonucleoprotein complex biogenesis 1.200000e-03
ribonucleoprotein complex biogenesis 4.700000e-04
ribosomal large subunit biogenesis 4.500000e-05
ribosome biogenesis 2.500000e-04
ribosome biogenesis 9.600000e-05
single organism membrane organization 2.000000e-17
single organism membrane organization 3.100000e-13
spliceosomal complex assembly 7.300000e-04
symbiosis encompassing mutualism through parasitism 4.600000e-12
symbiosis encompassing mutualism through parasitism 7.400000e-16
translation 3.000000e-18
translation 6.900000e-04
translation 7.400000e-14

translational elongation 2.800000e-26
translational elongation 7.800000e-20
translational initiation 2.400000e-18
translational initiation 2.800000e-24
translational termination 1.900000e-20
translational termination 3.900000e-27
viral gene expression 3.500000e-18
viral gene expression 4.700000e-24
viral latency 1.000000e-03
viral life cycle 1.900000e-21
viral life cycle 3.000000e-16
viral process 2.100000e-12
viral process 2.500000e-16
viral transcription 1.800000e-18
viral transcription 2.000000e-24

#### B.2 Gene Set: `go_cp_iea`

##### B.2.1 Gene Pathways Significant for WT- but not for AD-Biclusters

###### Choroid Plexus

cytosolic large ribosomal subunit 9.100000e-10
cytosolic part 5.600000e-18
cytosolic ribosome 3.700000e-21
cytosolic small ribosomal subunit 1.700000e-10
large ribosomal subunit 4.000000e-09
ribonucleoprotein complex 1.000000e-12
ribosomal subunit 1.800000e-19
ribosome 2.800000e-17
small ribosomal subunit 2.600000e-09

###### GABAergic Neurons

cytosolic large ribosomal subunit 2.600000e-20
cytosolic part 4.100000e-25
cytosolic ribosome 5.400000e-30
cytosolic small ribosomal subunit 2.700000e-10
large ribosomal subunit 1.700000e-18
mitochondrial proton transporting ATP synthase complex 2.700000e-03
proton transporting ATP synthase complex 3.500000e-03
ribonucleoprotein complex 6.400000e-18
ribosomal subunit 1.700000e-26
ribosome 5.000000e-22
small ribosomal subunit 1.200000e-08

##### Radial Glial Cells

apical part of cell 2.500000e-03
apical plasma membrane 1.400000e-03
basolateral plasma membrane 9.200000e-04
brush border 8.500000e-05
brush border membrane 3.300000e-05
cell projection membrane 1.300000e-03
melanosome 3.300000e-04
pigment granule 3.300000e-04

#### B.2.2 Gene Pathways Significant for AD- but not for WT-Biclusters

##### Astroglial Cells

U12 type spliceosomal complex 5.600000e-03
axon 4.000000e-03
catalytic step 2 spliceosome 4.900000e-07
cell body 1.300000e-04
clathrin sculpted vesicle 5.700000e-04
cytosolic large ribosomal subunit 1.400000e-47
cytosolic large ribosomal subunit 9.200000e-12
cytosolic part 1.500000e-04
cytosolic part 4.700000e-16
cytosolic part 5.300000e-63
cytosolic ribosome 4.800000e-06
cytosolic ribosome 5.800000e-82
cytosolic ribosome 7.100000e-20
cytosolic small ribosomal subunit 1.400000e-08
cytosolic small ribosomal subunit 1.500000e-04
cytosolic small ribosomal subunit 5.200000e-34
envelope 3.800000e-03
eukaryotic 43S preinitiation complex 1.800000e-02
eukaryotic 48S preinitiation complex 1.500000e-02
extracellular membrane bounded organelle 2.900000e-05
extracellular organelle 2.900000e-05
extracellular vesicular exosome 2.400000e-05
growth cone 1.800000e-03
large ribosomal subunit 7.600000e-11
large ribosomal subunit 9.300000e-42
melanosome 3.800000e-03
mitochondrial envelope 1.400000e-05
mitochondrial inner membrane 3.200000e-07
mitochondrial membrane 6.800000e-06
mitochondrial membrane part 7.600000e-12

mitochondrial part 5.100000e-04
mitochondrial proton transporting ATP synthase complex 1.000000e-07
mitochondrial respiratory chain 2.900000e-07
neuron part 1.900000e-05
neuron projection 1.600000e-04
neuronal cell body 8.200000e-05
organelle envelope 3.600000e-03
organelle inner membrane 1.500000e-06
oxidoreductase complex 9.800000e-04
phagocytic cup 1.500000e-02
pigment granule 3.800000e-03
polysome 5.600000e-03
proton transporting ATP synthase complex 2.400000e-07
proton transporting ATP synthase complex coupling factor F o 4.400000e-04
proton transporting two sector ATPase complex 4.500000e-06
proton transporting two sector ATPase complex proton transporting domain 3.700000e-03
respiratory chain 7.800000e-07
ribonucleoprotein complex 1.300000e-56
ribonucleoprotein complex 3.100000e-11
ribosomal subunit 2.200000e-71
ribosomal subunit 2.900000e-05
ribosomal subunit 7.500000e-18
ribosome 1.600000e-58
ribosome 3.000000e-04
ribosome 3.100000e-15
ruffle 2.900000e-03
site of polarized growth 1.900000e-03
small ribosomal subunit 1.300000e-28
small ribosomal subunit 2.000000e-07
small ribosomal subunit 7.200000e-04
spliceosomal complex 2.000000e-05
translation preinitiation complex 1.800000e-02

**Oligodendrocyte Progenitor Cells**

cytosolic large ribosomal subunit 2.400000e-05
cytosolic part 1.100000e-06
cytosolic ribosome 3.100000e-08
cytosolic small ribosomal subunit 8.100000e-04
extracellular membrane bounded organelle 9.800000e-05
extracellular organelle 9.800000e-05
extracellular vesicular exosome 9.000000e-05
large ribosomal subunit 5.600000e-05
ribonucleoprotein complex 1.100000e-06

ribosomal subunit 2.000000e-07
ribosome 2.300000e-06
small ribosomal subunit 2.300000e-03

**Ventral Progenitors**

U12 type spliceosomal complex 6.200000e-03
catalytic step 2 spliceosome 3.100000e-06
catalytic step 2 spliceosome 5.200000e-04
cell body 1.300000e-02
cell cortex 4.400000e-04
clathrin sculpted vesicle 1.700000e-03
cortical cytoskeleton 9.200000e-04
cytosolic large ribosomal subunit 1.900000e-16
cytosolic large ribosomal subunit 4.600000e-10
cytosolic part 1.800000e-21
cytosolic part 3.100000e-16
cytosolic ribosome 2.600000e-25
cytosolic ribosome 4.300000e-19
cytosolic small ribosomal subunit 2.500000e-08
cytosolic small ribosomal subunit 9.700000e-08
envelope 9.200000e-04
extracellular membrane bounded organelle 5.600000e-03
extracellular membrane bounded organelle 7.200000e-03
extracellular organelle 5.600000e-03
extracellular organelle 7.200000e-03
extracellular vesicular exosome 5.200000e-03
extracellular vesicular exosome 6.700000e-03
large ribosomal subunit 2.000000e-09
large ribosomal subunit 2.200000e-15
melanosome 1.400000e-03
mitochondrial envelope 8.800000e-05
mitochondrial inner membrane 4.500000e-03
mitochondrial inner membrane 7.200000e-03
mitochondrial membrane 3.300000e-04
mitochondrial membrane part 2.200000e-05
mitochondrial membrane part 4.100000e-04
mitochondrial part 4.300000e-04
mitochondrial proton transporting ATP synthase complex 3.000000e-03
mitochondrial respiratory chain 3.300000e-03
neuron projection 6.600000e-03
neuronal cell body 8.900000e-03
nuclear chromosome 1.100000e-02
nuclear chromosome telomeric region 5.200000e-03

nuclear chromosome part 5.800000e-03
nuclear speck 7.000000e-03
nuclear telomere cap complex 1.100000e-03
organelle envelope 8.700000e-04
organelle inner membrane 1.100000e-02
pigment granule 1.400000e-03
pronucleus 2.800000e-03
proton transporting ATP synthase complex 3.800000e-03
proton transporting two sector ATPase complex 1.400000e-03
respiratory chain 4.500000e-03
ribonucleoprotein complex 1.600000e-11
ribonucleoprotein complex 2.100000e-03
ribonucleoprotein complex 2.300000e-04
ribonucleoprotein complex 3.900000e-15
ribosomal subunit 1.400000e-17
ribosomal subunit 2.800000e-23
ribosome 1.300000e-15
ribosome 1.300000e-20
small ribosomal subunit 2.100000e-07
small ribosomal subunit 8.300000e-07
spliceosomal complex 5.800000e-04
spliceosomal complex 8.700000e-06
telomere cap complex 1.100000e-03

#### B.3 Gene Set: `go_mf_iea`

##### B.3.1 Gene Pathways Significant for WT- but not for AD-Biclusters

###### Choroid Plexus

RNA binding 3.000000e-10
structural constituent of ribosome 3.700000e-20
structural molecule activity 7.500000e-14

###### GABAergic Neurons

RNA binding 4.400000e-13
hydrogen ion transmembrane transporter activity 5.600000e-04
hydrogen transport 1.100000e-03
proton transport 1.100000e-03
rRNA binding 2.700000e-04
structural constituent of ribosome 3.800000e-26
structural molecule activity 1.600000e-18

**Radial Glial Cells**

GTP binding 2.500000e-05
guanyl nucleotide binding 2.800000e-05
guanyl ribonucleotide binding 2.800000e-05
ion channel binding 7.500000e-05
unfolded protein binding 1.800000e-04

##### B.3.2 Gene Pathways Significant for AD- but not for WT-Biclusters

**Astroglial Cells**

RNA binding 1.300000e-04
RNA binding 1.900000e-38
RNA binding 9.200000e-05
chaperone binding 1.800000e-04
cytochrome c oxidase activity 1.300000e-08
heme copper terminal oxidase activity 1.300000e-08
hydrogen ion transmembrane transporter activity 1.100000e-11
hydrogen transport 1.600000e-10
inorganic cation transmembrane transporter activity 1.800000e-03
mRNA binding 2.900000e-05
macromolecule transmembrane transporter activity 9.300000e-04
monovalent inorganic cation transmembrane transporter activity 3.100000e-05
monovalent inorganic cation transport 5.300000e-05
oxidoreductase activity acting on a heme group of donors 1.800000e-08
oxidoreductase activity acting on a heme group of donors oxygen as acceptor 1.300000e-08
protein transmembrane transporter activity 6.000000e-04
proton transport 1.600000e-10
rRNA binding 3.100000e-12
single stranded DNA binding 4.600000e-03
structural constituent of cytoskeleton 3.200000e-05
structural constituent of ribosome 1.100000e-17
structural constituent of ribosome 2.000000e-05
structural constituent of ribosome 8.000000e-69
structural molecule activity 1.300000e-11
structural molecule activity 1.900000e-43
syntaxin 1 binding 3.600000e-04
telomeric DNA binding 1.400000e-03
translation 2.600000e-07
translation elongation factor activity 1.800000e-04
translation factor activity nucleic acid binding 1.400000e-06
translation initiation factor activity 5.100000e-04
translational elongation 6.300000e-04
translational initiation 5.600000e-04

unfolded protein binding 3.700000e-06
unfolded protein binding 4.100000e-06

**Oligodendrocyte Progenitor Cells**

structural constituent of ribosome 1.600000e-07
structural molecule activity 9.700000e-06

**Ventral Progenitors**

GTP dependent protein binding 1.100000e-03
RNA binding 1.200000e-09
RNA binding 3.200000e-11
RNA binding 5.500000e-03
RNA binding 6.600000e-05
activation of cysteine type endopeptidase activity 2.600000e-03
activation of cysteine type endopeptidase activity involved in apoptotic process 2.600000e-03
cation transmembrane transporter activity 2.400000e-03
cation transport 2.500000e-03
chaperone binding 9.200000e-04
cysteine type endopeptidase activator activity involved in apoptotic process 2.600000e-03
cytochrome c oxidase activity 2.000000e-06
establishment of protein localization 3.900000e-03
heme copper terminal oxidase activity 2.000000e-06
hydrogen ion transmembrane transporter activity 1.600000e-06
hydrogen transport 4.900000e-06
inorganic cation transmembrane transporter activity 6.300000e-04
low density lipoprotein particle receptor binding 1.500000e-03
mRNA binding 3.500000e-05
macromolecule transmembrane transporter activity 2.800000e-05
monovalent inorganic cation transmembrane transporter activity 2.000000e-03
monovalent inorganic cation transport 2.600000e-03
oxidoreductase activity acting on a heme group of donors 2.400000e-06
oxidoreductase activity acting on a heme group of donors oxygen as acceptor 2.000000e-06
peptidase activator activity involved in apoptotic process 3.200000e-03
positive regulation of cysteine type endopeptidase activity 2.600000e-03
positive regulation of cysteine type endopeptidase activity involved in apoptotic process 2.600000e-03
positive regulation of endopeptidase activity 4.200000e-03
posttranscriptional regulation of gene expression 6.100000e-03
protein localization 3.900000e-03
protein transmembrane transporter activity 1.800000e-05
protein transport 3.900000e-03
protein transporter activity 3.900000e-03
proton transport 4.900000e-06
regulation of translation 6.100000e-03

single organism transport 1.700000e-03
structural constituent of ribosome 3.400000e-18
structural constituent of ribosome 4.400000e-24
structural molecule activity 1.500000e-12
structural molecule activity 1.700000e-16
substrate specific transmembrane transporter activity 3.800000e-04
substrate specific transporter activity 1.200000e-03
syntaxin 1 binding 1.300000e-03
thioesterase binding 1.300000e-03
translation 1.100000e-03
translation 1.600000e-03
translation factor activity nucleic acid binding 5.100000e-04
translation initiation factor activity 1.400000e-03
translation regulator activity 3.200000e-03
translational initiation 1.500000e-03
transmembrane transport 8.600000e-04
transmembrane transporter activity 8.600000e-04
unfolded protein binding 3.800000e-05
unfolded protein binding 8.100000e-04
